# Spatial patterning of fibroblast TGFβ signaling underlies treatment resistance in rheumatoid arthritis

**DOI:** 10.1101/2025.03.14.642821

**Authors:** Kartik Bhamidipati, Alexa B R McIntyre, Shideh Kazerounian, Gao Ce, Miles Tran, Sean A Prell, Rachel Lau, Vikram Khedgikar, Christopher Altmann, Annabelle Small, Vincent Wong, Roopa Madhu, Sonia Presti, Ksenia S Anufrieva, Philip E Blazar, Jeffrey K Lange, Jennifer Seifert, Accelerating Medicines Partnership: RA/SLE Network, Accelerating Medicines Partnership: Autoimmune and Immune-Mediated Diseases Network (AMP-AIM), Larry W Moreland, Adam P Croft, Myles J Lewis, Ranjeny Thomas, Anna H Jonsson, Costantino Pitzalis, Ellen M Gravallese, Michael B Brenner, Ilya Korsunsky, Mihir D Wechalekar, Kevin Wei

## Abstract

Treatment-refractory rheumatoid arthritis (RA) is a major unmet need, and the mechanisms driving treatment resistance are poorly understood. To identify molecular determinants of RA non-remission, we performed spatial transcriptomic profiling on pre- and post-treatment synovial tissue biopsies from treatment naïve patients who received conventional DMARDs or adalimumab for 6 months. In the baseline biopsies of non-remission patients, we identified significant expansion of fibrogenic fibroblasts marked by high expression of *COMP*, a fibrosis-associated extracellular matrix protein. *COMP*hi fibroblasts localized to perivascular niches that, unexpectedly, served as transcriptional hubs for TGFβ activity. We identified endothelial-derived Notch signaling as an upstream regulator of fibroblast TGFβ signaling via its dual role in driving TGFβ isoform expression and suppressing TGFβ receptors, generating a proximal-distal gradient of TGFβ activity. Further, disruption of steady-state Notch signaling *in vitro* enabled fibrogenic fibroblast activation. Analysis of post-treatment biopsies revealed marked expansion of *COMP*hi fibroblasts in non-remission RA patients, despite evidence of successful immune cell depletion, suggesting a spatiotemporal process of fibrogenic remodeling linked to treatment resistance. Collectively, our data implicates targeting of TGFβ signaling to prevent exuberant synovial tissue fibrosis as a potential therapeutic strategy for refractory RA.

## Introduction

Rheumatoid arthritis (RA) is a common autoimmune disease characterized by chronic inflammation in the synovium^1,2^. While there have been significant advances in the treatment of RA with the introduction of biologics targeting pathogenic inflammatory mediators, treatment-refractory RA remains a major challenge^3^. Greater than 50% of patients do not achieve remission with initial lines of therapy and 5-30% of patients remain unresponsive to multiple lines of therapy^4,5^. Clinically, such treatment refractory patients present more frequently with concomitant non-inflammatory pain, suggesting an alternate pathophysiology^6^. However, the cellular and molecular mechanisms underlying refractory RA remain poorly understood. Synovial fibroblasts are mesenchymal cells that with macrophages constitute the joint lining membrane^7^. In RA, fibroblasts undergo expansion and acquire pathological states that sustain inflammation^8,9^ ^9–11^ and drive joint damage^10^. Transcriptomic analysis of RA synovium has revealed high phenotypic and functional diversity among synovial fibroblasts^8,9,11–14^ and fibroblastic gene signatures that predict treatment failure^15–17^.

Fibrosis is a pathogenic process characterized by exuberant fibroblast activation, leading to the accumulation of connective tissue components and extracellular matrix (ECM)^18^ with TGFβ as a central mediator of this process^19^. Fibrosis leading to organ damage is typically associated with autoimmune diseases such as systemic sclerosis and interstitial lung disease and is a well-established contributor to joint stiffness and pain in osteoarthritis (OA)^20^. RA, in contrast, is considered a chronic inflammatory disease; therefore, the potential contributions of synovial tissue fibrosis to RA treatment response and its mechanistic origins early in disease have not been well studied.

We have previously uncovered the role of spatial context in driving signals that generate fibroblast diversity and define positional identity^9,21^. Until recently, spatially aware transcriptomic profiling of fibroblasts required multiple iterations of protein or RNA detection, which poses a major barrier to high-dimensional characterization of cellular niches and associated signaling. Here, we applied a subcellular resolution, high-dimensional (5,101 genes) spatial transcriptomic technology to pre- and 6-month post-treatment synovial biopsies to identify spatial determinants of treatment resistance in recent-onset RA. At this stage, there is a window of opportunity to achieve remission, which predicts better outcomes in RA^22^. In the baseline biopsies of patients not achieving remission, we identified significant expansion of fibrogenic fibroblasts, marked by high expression of *COMP*. *COMP*hi fibroblasts localized to perivascular niches where we observed coordinated spatial patterning of TGFβ activity. Mechanistically, endothelial-derived Notch signaling actively finetunes the TGFβ-mediated fibrogenic program by simultaneously inducing TGFβ expression and suppressing TGFβ receptor expression, resulting in the establishment of an endothelial proximal-to-distal transcriptional gradient among fibroblasts. Perturbation of steady-state Notch signaling *in vitro* resulted in the disruption of the proximal-to-distal fibroblast axis, leading to the de-repression of fibrogenic gene expression. In post-treatment biopsies, we observed significant expansion of *COMP*hi fibroblasts in non-remission patients, despite evidence of successful immune depletion. Together, these data suggest that TGFβ signaling and fibrogenic fibroblast activation drive a treatment-refractory tissue phenotype and that this process can be captured at the earliest stages of RA. Targeting TGFβ signaling could therefore represent an additional strategy to address RA treatment resistance.

## Results

### Identification of distinct immune and fibroblastic niches in RA synovium

To identify spatial features associated with non-remission, we performed spatial transcriptomics analysis on 17 treatment-naïve biopsies from patients with early RA as part of the 396.10 study (**Fig. 1a**). Newly diagnosed patients were randomly assigned to triple therapy disease-modifying antirheumatic drugs (DMARDs) or targeted therapy (TNFi). Remission status, as defined by DAS28-ESR < 2.6, were recorded after six months of treatment with either anti-TNF biologic adalimumab or triple therapy conventional synthetic DMARD (methotrexate, sulfasalazine, and hydroxychloroquine) (**Supplementary Table 1**). We first used a cell type-naïve clustering approach that leverages spatial differences in transcript expression to draw boundaries between tissue compartments and identify major synovial tissue niches (**Fig. 1b, Extended Data Fig. 1a**)^23^. After excluding one sample based on transcript sparsity, we identified seven niches which were enriched for distinct cell types in treatment-naïve biopsies: 1) fibroblast-rich 2) vascular, 3) stromal-adipose, 4) lining fibroblasts (“liningF”), 5) lining macrophages (“liningM”), 6) T and B cells (“immuneT”), and 7) plasma cells (“immuneP”) (**Fig. 1c-d**, **Extended Data Fig. 1a-b**). The lining niches (liningF, liningM) consisted of a mix of synovial lining fibroblasts (*CD55+*) and lining macrophages (*HTRA1*+). The immune niches contained dense infiltrates composed of either primarily T cells (*CD3E+*) and B cells (*MS4A1*+) (“immuneT”) or plasma cell aggregates (*MZB1*+) (“immuneP”). Vascular niches consisted of various endothelial subtypes including venules (*SELP*+), capillaries (*PLVAP*+), and arterioles (*PODXL*+), mural cells, including pericytes (*RGS5*+) and vascular smooth cells (*ACTA2*+), and *NOTCH3*+ vascular fibroblasts. Stromal-adipose niches contained vascular cells and fibroblasts embedded among adipocytes (*ADIPOQ*+). Fibroblast-rich niches were composed primarily of sublining fibroblasts (*COL5A1*hi).

**Figure 1.**
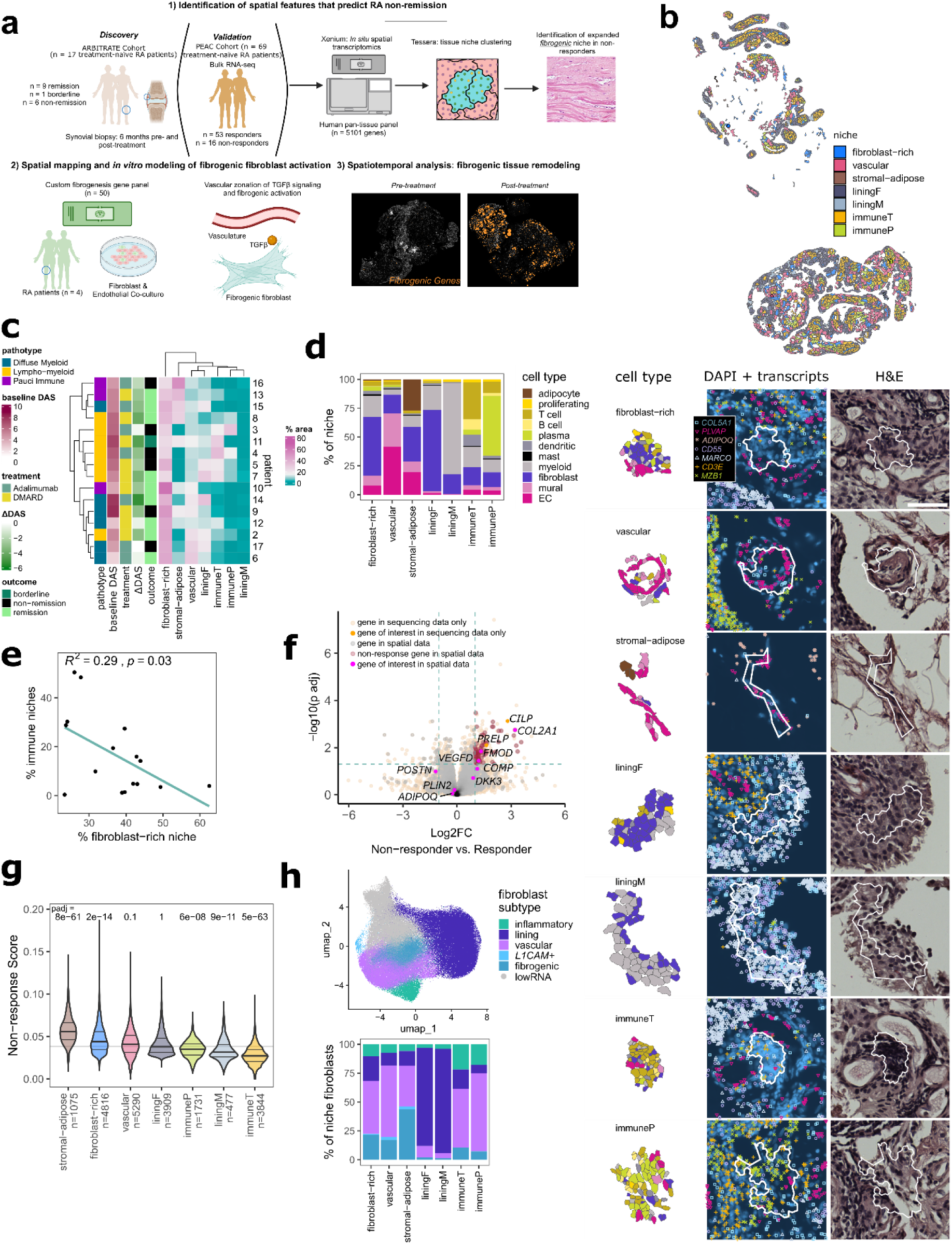
Identification of fibroblastic and immune niches in the treatment-naïve RA synovium. **a,** Schematic representing an overview of the study. **b,** Visualization of niches in a single synovial biopsy. Representative examples for each individual niche type with component cell types labelled are shown below with corresponding DAPI and H&E images. Scale bar indicates 50 µm. **c,** Heatmap representing the abundance of tissue niches, by tissue area, in each treatment naïve biopsy and the associated clinical metadata. Baseline DAS and ΔDAS represent the DAS28 ESR pre-treatment and the change at 6-months, respectively. **d,** Stacked bar plot representing the proportion of cell types per niche. **e,** Correlation plot showing the relationship between relative area of immune niches (immuneT and immuneP) and the total area of fibroblast-rich niches per sample. **f,** Volcano plot representing differentially expressed genes in non-responders versus responder from bulk RNA-sequencing analysis of synovial tissue derived from treatment naïve patients. Selected genes, including those in the non-responder gene set, are highlighted. **g,** Violin plot representing the distribution of non-response scores per niche type, with the number of niches analyzed indicated. Statistical comparison by two-sided Mann-Whitney U comparing a down-sampled selection of non-response scores in 500 niches from each niche type to a random selection of scores from 500 other niches. Horizontal line represents the median non-response score across all niches analyzed. **h,** Uniform Manifold Approximation and Projection (UMAP) plot of fibroblasts annotated by subtype, and stacked bar plot representing the proportion of each subtype within the fibroblast compartment of each niche.

The relative composition of the seven niches markedly varied across patients (**Fig. 1c-d, Extended Data Fig. 1c)**; hierarchical clustering based on the tissue area covered by niches classified samples into three broad categories, one characterized by stromal-adipose enrichment (n = 4), another by immune niche enrichment (n = 5) and a third marked by expanded vascular and fibroblast niches (n = 7) (**Fig. 1c**.) While we observed high concordance between immune niche tissue area and H&E-based lympho-myeloid pathotype classification^24^, (p = 0.007 and p = 0.0033 in comparisons to diffuse myeloid and pauci-immune samples, two-sided Mann-Whitney U-tests with Bonferroni correction) (**Extended data Fig. 1e**), clear distinctions in niches did not appear between the diffuse myeloid and pauci-immune pathotypes, suggesting that spatial transcriptomics analysis captures complexity in cellular organization beyond traditional histologic assessment of synovial tissue pathotype.

At a niche level, the tissue area of fibroblast-rich niches negatively correlated with the area of combined immune niches (R = −0.54, p = 0.03, two-sided Pearson correlation coefficient test) (**Fig.1e, Extended data Fig. 1d**), suggesting that the key determinant of RA synovial tissue heterogeneity is the abundance of immune-cell infiltrates versus expansion of fibroblastic regions, consistent with the clustering of immuneT and immuneP niches in biopsies with lympho-myeloid pathotype (**Fig. 1c**) and prior studies utilizing histology^25,26^ or single-cell transcriptomics^12,15,16^. When we evaluated the relationship between sample-level niche composition at baseline and remission status at 6 months, we observed no statistically significant differences between groups, suggesting that broad tissue composition alone was not sufficient to stratify patients based on remission status (**Extended Data Fig. 1f**).

### *COMP* expression defines a fibrogenic program associated with non-remission

Since the sample size of our cohort is limited, we leveraged bulk RNA-sequencing data from the larger PEAC cohort (n = 69 treatment-naïve patients, with n = 53 responders and 16 non-responders defined by DAS28-CRP > 5.1 and DAS28-CRP improvement ≤ 0.6 after 6 months) ^16^ to identify genes associated with non-response (log2 fold change > 1, padj < 0.05 between non-responders and responders, n = 290 genes) (**Fig. 1f**). We applied the non-responder gene signature (n = 73 genes that overlapped with the spatial transcriptomics panel), to spatial niches and found enrichment of the signature in stromal-adipose (padj = 8e-61, two-sided Mann-Whitney U test compared to randomly selected niches), fibroblast-rich (padj = 2e-14) and vascular niches (padj = 0.1) compared to the immune and lining niches (**Fig. 1g**). Among major synovial cell types, fibroblasts exhibited the highest non-response scores (**Extended Data Fig. 1g**).

To identify specific fibroblast phenotypes most strongly associated with non-response, we subtyped fibroblasts^15^ in our spatial transcriptomic data and identified four sublining subsets in addition to lining, including fibrogenic, characterized by high extracellular matrix gene (ECM) gene expression (*COL6A1*, *COL8A1*, *COMP*), inflammatory, characterized by expression of inflammatory mediators (*CXCL12, SFRP1)*, vascular, characterized by *NOTCH3* and *THY1* expression, and a minor subpopulation marked by high *L1CAM* (**Fig. 1h, Extended Data Fig. 1h**). We observed a fifth cluster of sublining fibroblasts (lowRNA) that was difficult to classify given low overall detection of transcripts and defining markers. Examining the fibroblast compartment of each niche, we confirmed that lining fibroblasts were most abundant in the lining niches and that vascular and inflammatory fibroblasts were most enriched in the immune niches, consistent with the presence of immune aggregates around postcapillary venules^27^. Notably, fibrogenic fibroblasts were specifically expanded in the non-responder associated fibroblast-rich, stromal-adipose and vascular niches (**Fig. 1h**) and were also the subtype most enriched for the non-responder gene signature (**Extended Data Fig. 1i.)**

While the vascular and inflammatory fibroblast compartments have previously been characterized^8–11,13^, the fibrogenic compartment in RA remains incompletely understood^25^. To better characterize fibrogenic fibroblasts in RA and their potential contribution to non-remission, we first defined a 10-gene fibrogenic signature based on genes upregulated in disease-associated fibroblast populations, annotated in previously published systemic sclerosis (SSc) and idiopathic pulmonary fibrosis (IPF) datasets^28,29^ as scleroderma associated fibroblasts (ScAFs) and myofibroblasts respectively (**Extended data Fig. 2a**). The gene signature consisted of fibrosis-associated collagens such as *COL1A1* and *COL3A1* as well as other fibrosis-associated ECM components including *SPARC* and *ASPN*^30,31^ ^32^. We then subsetted and re-clustered fibroblasts from patients (n = 82) in the AMP RA/SLE single-cell RNA sequencing dataset^15^ that were highly enriched for the fibrogenic gene signature (top 10% of cells) and observed two broad fibrogenic fibroblast clusters – one marked by high *COMP* expression and another marked by high *POSTN* expression (**Fig. 2a-b)**. Interestingly, while ECM genes were highly expressed by both fibroblast subsets, *POSTN*hi fibroblasts had relatively higher expression of collagen genes (*COL1A1*, *COL3A1*, *COL8A1*) whereas the *COMP*hi subset more strongly expressed genes previously found to be associated with treatment resistance (*DKK3, FMOD, PRELP)*^17^, illustrating distinct phenotypes among fibrogenic fibroblast (**Fig. 2b)***. COMP*hi fibroblasts exhibited a striking enrichment of the non-response signature compared to the *POSTN*hi subset (p < 2.22e-16), suggesting that *COMP* specifically marks a subset of fibrogenic fibroblasts associated with treatment non-response (**Fig. 2c)**.

**Figure 2.**
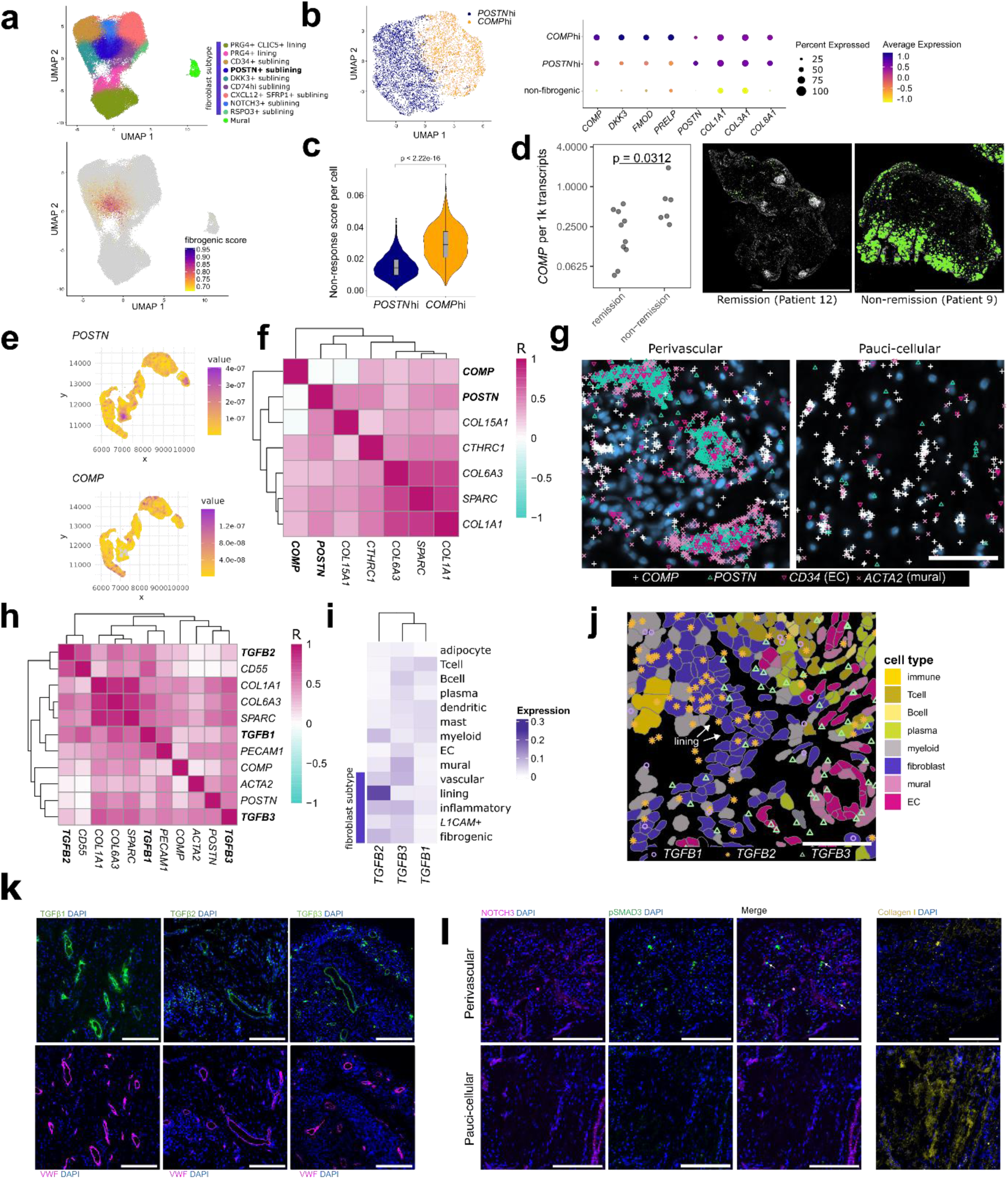
*COMP* and *POSTN* define spatially segregated fibrogenic subsets that co-localize with TGFβ activity. **a,** UMAP projection of synovial fibroblasts with cluster annotations (ref.^15^) and projection of the fibrogenic gene signature score onto the UMAP as calculated by UCell. Fibroblasts with signature scores in the top 10% (>0.6713282) are colored. **b**, UMAP projection of subsetted and re-clustered fibroblasts that had fibrogenic signature scores in the top decile. Dotplot of selected genes differentially expressed across clusters with the color representing scaled expression and size of the dot representing percent of cells expressing the gene. Non-fibrogenic refers fibroblasts from the full synovial dataset that had fibrogenic signature scores in the bottom 90 percentile. **c,** Violin plot displaying the distribution of the non-responder signatures (n = 290 genes) in *COMP*hi and *POSTN*hi cells, as calculated by Ucell. Wilcoxon test was used for statistical comparisons between the groups**. d,** Quantification of normalized *COMP* transcript expression in baseline non-remission and remission biopsies, with representative images of *COMP* transcript expression. Scale bar indicates 2 mm **e,** Visualization of kernel density estimates showing correlation between *POSTN* and *COMP* expression within a representative *COL1A1*-high region. **f,** Heatmap showing mean pixel correlations across synovial samples (n = 4) between kernel densities for selected fibrogenic genes within *COL1A1*-high regions. **g,** Representative examples of *COMP* transcript expression in perivascular and pauci-cellular regions. Scale bar indicates 50 µm. **h,** Mean pixel correlations across samples (n = 4) for *TGFB* isoform expression compared to selected fibroblast and endothelial markers. **i,** Heatmap representing relative expression of *TGFB* isoforms across different cell types synovial samples (n = 17) analyzed with a 5101-gene panel. **j,** Example of data in (i) showing *TGFB* transcripts overlaid on cell types. Scale bar indicates 50 µm. **k,** Immunofluorescence staining of RA synovial tissue showing protein expression of TGFβ isoforms relative to endothelial cell marker VWF. TGFβ3 and VWF staining were on serial sections. **l,** IF data showing NOTCH3, Collagen 1, and pSMAD3 staining in perivascular and pauci-cellular regions of the RA synovium. Scale bars are 200 µm.

To test whether fibrogenic gene expression could stratify remission and non-remission patients, we examined the expression of fibrogenic transcripts in the 396.10 cohort. We found that *COMP*, but not *DKK3*, *FMOD* or *POSTN,* was significantly elevated in the treatment-naïve biopsies of patients who failed to achieve remission (*COMP*: p = 0.0312, *DKK3*: p = 0.87, *FMOD*: p = 0.43, *POSTN*: p = 0.428, two-sided Mann-Whitney U tests, **Fig. 2d, Extended Data Fig. 2b**). Collectively, differential transcript expression in our cohort supports *COMP*hi fibroblast abundance as a marker of treatment resistance.

### Perivascular compartmentalization of TGFβ signaling

To localize *COMP*hi fibroblasts and better characterize synovial tissue fibrosis, we designed a targeted panel of 50 fibrogenesis-associated genes for spatial transcriptomics analysis of additional RA synovial tissue samples (n = 4). The smaller panel enhanced the detection of relevant transcripts, including TGFβ isoforms, key regulators of fibrosis^19^, and *COMP* (mean *COMP* transcripts per cell = 2.4 in the targeted 50-gene panel vs. 0.17 in 5k data)^33,34^. Within fibroblast-rich tissue regions, defined by high *COL1A1* expression, we found that *COMP* and *POSTN* transcript kernel density estimates mapped to spatially distinct areas (**Fig. 2e-f**), supporting the reciprocal expression pattern observed in single-cell synovial data. We observed two major patterns of *COMP*hi fibroblast localization: 1) within pauci-cellular fibroblast-rich regions and 2) within the distal regions of perivascular niches, in the layers surrounding mural cells and vascular fibroblasts (**Figure 2g**). Accordingly, patient-level analysis of the spatial data revealed a positive association between the proportion of vascular cells and the abundance of *COMP*-expressing fibroblasts in synovial data (custom panel: R^2 = 0.66, p = 0.19, two-sided Pearson’s correlation coefficient test, 396.10: R^2 = 0.16, p = 0.12, two-sided Pearson’s correlation coefficient test) (**Extended Data Fig. 2c**). Additionally, we observed an unexpectedly strong association between *POSTN* transcripts and mural cell (*ACTA2*) and vascular fibroblast (*COL15A1*) markers, indicating strong expression of *POSTN* in the proximal cell layers of perivascular niches. (**Fig. 2h, Extended Data Fig. 2d**). Together, our data indicates that both subsets of fibrogenic fibroblasts localize to the perivascular compartment in RA synovia.

We next asked whether TGFβ expression is spatially associated with fibrogenic niches in RA by examining the spatial patterning of the TGFβ isoforms (**Fig. 2h**). *TGFB1* co-localized with the lining fibroblast marker *CD55* and endothelial cell marker *PECAM1*. *TGFB2* transcripts strongly colocalized with lining transcripts (*CD55*). *TGFB3* transcripts colocalized not only with markers of endothelial (*PECAM1*) and mural (*ACTA2*) cells, suggesting vascular and perivascular enrichment, but also overlapped with fibrogenic transcripts, including subpopulations marked by *POSTN*, *COMP* and other ECM genes *COL1A1, COL6A3, and SPARC*.

We examined TGFβ expression in additional cell types using data from the 396.10 cohort Xenium 5k data (**Fig. 2i-j, Extended Data Fig. 2e**). *TGFB1* showed low transcript detection relative to the other isoforms in this dataset but was expressed by myeloid and immune populations. *TGFB2* showed the highest expression in lining fibroblasts and myeloid cells. *TGFB3* was stromal-enriched and, consistent with the density analysis, was strongly expressed in the vascular compartment, including in vascular fibroblasts and mural cells.

Immunofluorescent staining of synovial tissue confirmed perivascular localization of TGFβ1, TGFβ2, and TGFβ3 at the protein level (**Figure 2k**), though TGFβ2 also showed high expression in immune aggregates, potentially bound to its receptor. Accordingly, we observed that a specific indicator of active TGFβ signaling, pSMAD3, was restricted to perivascular niches and absent in pauci-cellular regions characterized by more dense collagen deposition (**Fig. 2l, Extended Data Fig. 2f**). This suggests that active TGFβ signaling occurs specifically within fibroblasts in the perivascular compartment. Together, the observed spatial patterning of both fibrogenic gene expression and TGFβ suggests that the perivascular niche is an important hub for fibrogenic signaling.

### Endothelial cells generate spatial patterning of TGFβ responsiveness in fibroblasts

To better understand how vascular endothelial cells create spatial patterning of stromal TGFβ signaling, we modeled endothelial–fibroblast interactions *in vitro* by co-culturing human umbilical vein endothelial cells (HUVEC) with synovial fibroblasts (**Fig. 3a-b**). We spatially profiled this system using the custom fibrogenic gene panel. We observed distinct fibroblast transcriptional programs as a function of proximity to the nearest endothelial cell: fibroblasts in the most proximal layer expressed Notch target genes including *JAG1*, *HES1* and *NOTCH3*, which decreased in distal layers of fibroblasts that were more than approximately 1 cell layer away (median intercellular distance of ∼23 µm) (**Fig. 3c-e**) (p < 2.22e-308, two-sided Mann-Whitney U tests comparing distances from ECs for the top vs. bottom quantiles of cells in expression of each gene). Like Notch target genes, *POSTN* expression was also highest on the proximal layer of fibroblasts adjacent to endothelial cells and gradually decreased in fibroblasts distal from ECs (p = 5.27e-33, two-sided Mann-Whitney U test). In contrast, *COMP* expression was lowest in the endothelial-proximal layer but increased in distal fibroblasts (p = 1.36e-111, two-sided Mann-Whitney U test) (**Fig. 3f-h**). Overall, our endothelial-fibroblast model suggests that endothelial cell-derived signals pattern fibrogenic gene expression in a proximal to distal pattern where *POSTN* represents an EC-proximal fibrogenic pattern and *COMP* expression represents an EC-distal fibrogenic pattern.

**Figure 3.**
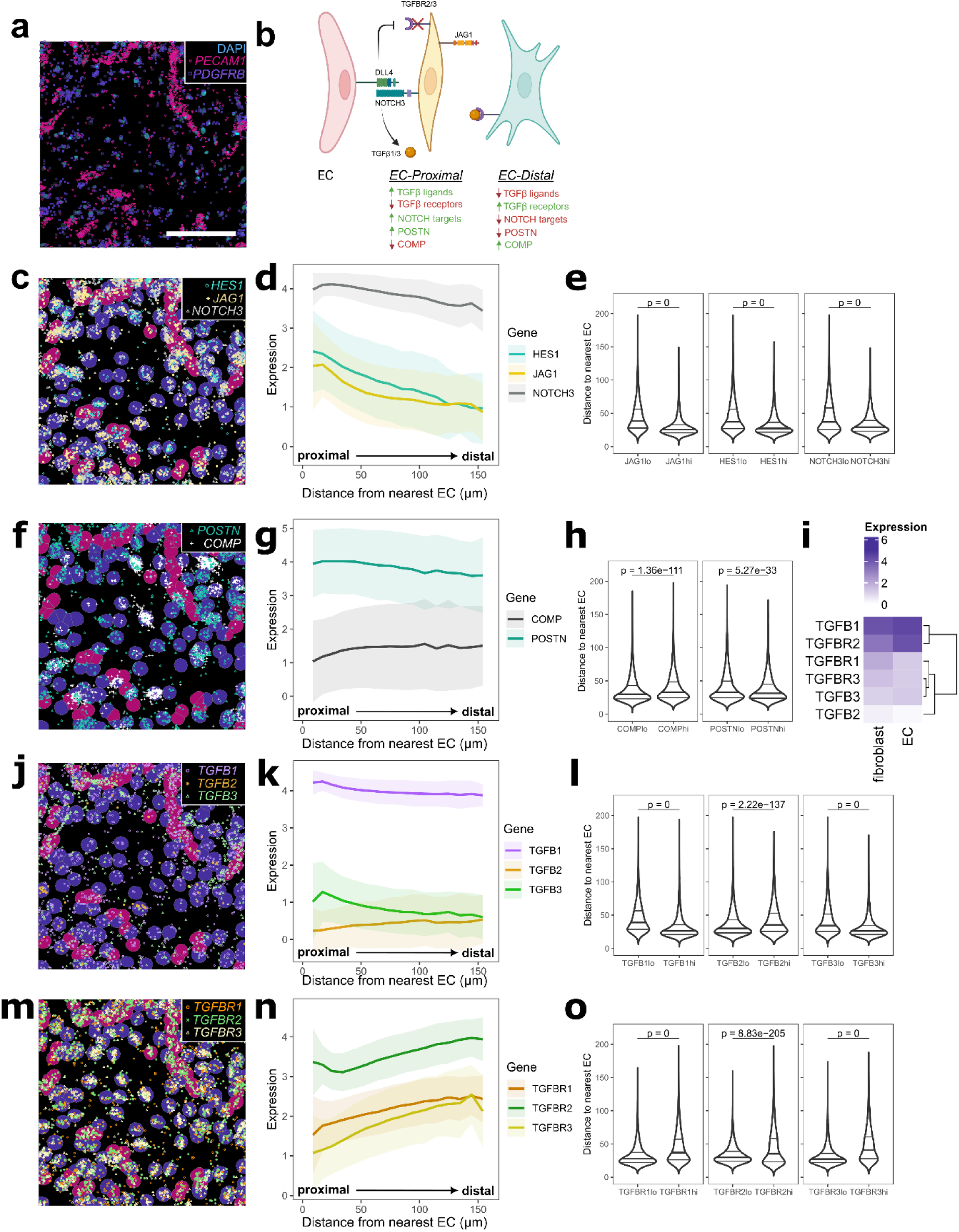
Endothelial cells generate a proximal to distal gene expression pattern in surrounding fibroblasts. **a,** Representative visualization of spatially profiled co-culture with transcripts for fibroblast marker, *PDGFRB*, and endothelial marker, *PECAM1*, overlaid on DAPI. Scale bar indicates 200 µm. **b,** Schematic summarizing the spatial distribution of fibroblast transcripts based on endothelial cell proximity. **c,f,j,m,** Example images of gene expression distribution overlaid on labelled cell types, to a maximum of randomly selected 1500 transcripts per gene, with Notch target genes shown in (c), fibrogenic markers in (f), *TGFB* ligands in (j), and *TGFB* receptors in (m). **d,g,k,n,** Line plot representing average expression of transcripts per fibroblast relative to the EC distance, with Notch target genes shown in (d), fibrogenic markers in (g), *TGFB* ligands in (k), and *TGFB* receptors in (n). Solid line shows mean, shaded regions show one standard deviation. **e,h,l,o,** Distance to the nearest EC in the top vs. bottom quantile of cells by expression of each gene, with Notch target genes shown in (e), fibrogenic markers in (h), *TGFB* ligands in (l), and *TGFB* receptors in (o). **i,** Heatmap representing expression of *TGFB* ligands and receptors by cell type in the co-culture.

Next, we evaluated the distribution of TGFβ isoforms along the EC proximal-distal axis. Consistent with the expression pattern in RA synovia, *TGFB1* was expressed on endothelial cells and on the EC-proximal fibroblast layer, with fewer average transcripts in EC-distal fibroblasts (p < 2.22e-308, two-sided Mann-Whitney U test) (**Fig. 3i-l**). *TGFB2* was lowly expressed and showed an increasing trend with distance from endothelial cells (p = 2.22e-137, two-sided Mann-Whitney U test). *TGFB3*, like *TGFB1,* was induced on endothelial-proximal fibroblasts and then sharply decreased in EC-distal fibroblasts (p < 2.22e-308, two-sided Mann-Whitney U test). We also measured the spatial distribution of fibroblast TGFβ receptor expression to nominate receptor-ligand pairs responsible for generating differential fibroblast TGFβ response. Transcripts of *TGFBR1*, the TGFβ signaling co-receptor, *TGFBR2*, the ligand binding receptor, and *TGFBR3*, a TGFβ binding co-receptor^35^, were lowest in EC-proximal layers and gradually increased in EC-distal fibroblasts (p < 2.22e-308, p = 8.83e-205, p < 2.22e-308 respectively, two-sided Mann-Whitney U tests) (**Figure 3m-o**). Collectively, these data suggest that endothelial signals determine spatial patterning of TGFβ signaling via paired regulation of TGFβ ligands and receptors.

### Patterning of fibroblast TGFβ responsiveness via Notch signaling

Given the distinct spatial pattern of TGFβ-related transcripts in fibroblasts generated by contact with neighboring endothelial cells, we evaluated the potential role of endothelial-derived Notch in regulating TGFβ signaling. We re-analyzed scRNAseq data from matrigel-embedded 3D organoids of fibroblasts and ECs treated with or without the gamma-secretase inhibitor DAPT, which inhibits Notch signaling^9^ (**Fig. 4a-b**,). We observed significant induction of *TGFB1* (log2FC 2.08539, padj 0, Wilcoxon Test), *TGFB3* (log2FC 3.054497, padj 1.233-118) and fibrogenic gene expression (*COL1A1*: log2FC 4.974715, padj 0) in fibroblast-EC organoids as well as suppression of *TGFBR2 (*log2FC −0.8789, padj 2.800e-115) and *TGFBR3* (log2FC −2.06822, padj 2.842e-284) expression, which was strongly reversed with the addition of DAPT, directly implicating Notch signaling as an upstream regulator of fibroblast TGFβ signaling (**Fig. 4c-d, Extended Data Fig. 3a**).

**Figure 4.**
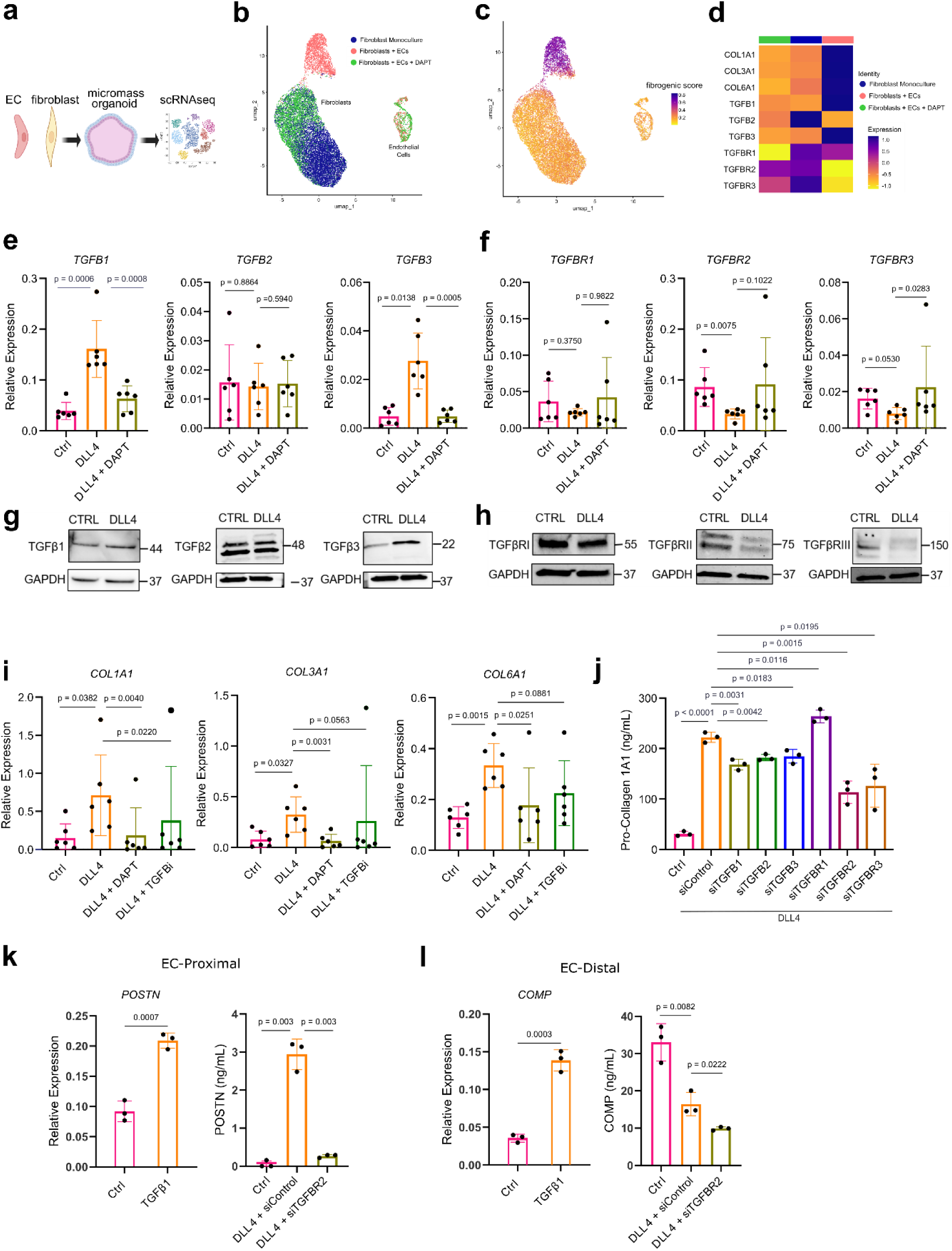
Notch signaling controls TGFβ isoform and receptor expression in synovial fibroblasts. **a-b,** Schematic of experimental workflow and UMAP plot of isolated single cells from the indicated micromass organoid conditions (ref^9^). Fibroblast Monoculture: organoid containing fibroblasts only; Fibroblasts + ECs: organoids containing fibroblasts and ECs; Fibroblasts + ECs + DAPT: organoids containing fibroblasts and ECs treated with a NOTCH inhibitor. **c,** UMAP plot of cells from organoid culture shaded by level of fibrogenic gene signature score as calculated with UCell. **d,** Heatmap of selected genes grouped by experimental condition. **e-f,** RT-qPCR analysis of *TGFB* isoform and receptor gene expression on unstimulated or DLL4 stimulated fibroblasts treated with or without DAPT (10 uM) for 72 hours. **g-h,** Immunoblots of TGFβ isoforms and receptors with lysates from unstimulated or DLL4 stimulated fibroblasts (72h). **i,** RT-qPCR analysis of *COL1A1*, *COL3A1* and *COL6A1* gene expression on unstimulated or DLL4 stimulated fibroblasts treated with or without TGFβ inhibitor (SB431542; 10 uM) or DAPT (10 uM) for 72h. **j**, ELISA quantification of fibroblast production of pro-collagen I alpha 1 over 24h after 5 days of treatment with siRNA (20nM) during or without DLL4 stimulation. **k-l**, RT-qPCR and ELISA quantification of POSTN (k) and COMP (l) after 72 hours of stimulation with recombinant TGFβ1 (10 ng ml−1) (left) or with DLL4 stimulation and siRNA targeting of TGFBR2 (right). For **e,f,i** each data point represents an independent cell line (n = 6) and for **j-l** each data point represents biological replicates (n = 3) from a single cell line and are representative of at least two independent experiments. Data are shown as mean +/- s.d. Statistical analysis was performed by a two-sided ratio paired t-test for **e**, **f, i** and unpaired two-sided Student’s t-test for **j-l.**

To directly examine the effects of fibroblast Notch activation on TGFβ signaling, we stimulated synovial fibroblast cell lines (n = 6) with plate-bound DLL4 and observed strong induction of *TGFB1 (*4.121 fold change, p = 0.0006) and *TGFB3* (5.646 fold change, p = 0.0138) gene expression that was reversed with DAPT (**Fig. 4e-f**). We also observed strong suppression of *TGFBR2* (−2.613 fold change, p = 0.0075) and moderate suppression of *TGFBR3* (−2.006 fold change, p = 0.0530) that was significantly reversed with the addition of gamma secretase-inhibitor DAPT (p = 0.0283). We confirmed protein-level induction of fibroblast TGFβ1 and TGFβ3 and suppression of TGFβRII and TGFβRIII in response to DLL4 stimulation (**Fig. 4h)**. Further supporting Notch receptor signaling as an upstream regulator of TGFβ signaling, we analyzed publicly available ChIP-seq data for binding of the Notch-associated transcription factor, recombination signal binding protein for immunoglobulin kappa J region (RBPJ), in HepG2 cells and observed significant peaks in the promoter regions of *TGFBR2* (−350 bp to −138 bp upstream of transcriptional start site (TSS)) and *TGFBR3* (−185 bp to +84 bp relative to TSS), supporting a role for Notch in direct transcriptional regulation of TGFβ receptors (**Extended Data Fig. 4b-c**).

Next, we assessed the impact of Notch and TGFβ signaling on fibrogenic gene expression downstream of DLL4 stimulation and observed strong induction of fibrogenic collagen expression, including *COL1A1* (4.75 fold change, p = 0.0392), *COL3A1* (4.00 fold change, p = 0.0327) and *COL6A1* (2.60 fold change, p = 0.0015) that was reversed with the addition of DAPT or TGFβ inhibitor (SB-431542) (**Fig. 4i**). We then systematically tested the requirement of each individual TGFβ isoform and TGFβ receptor in the Notch-activated fibrogenic program (**Fig. 4j**). DLL4-stimulated fibroblasts produced high amounts of pro-collagen I alpha, compared to unstimulated fibroblasts (222.2 ng/mL vs 30.92 ng/mL, p <0.0001), which was significantly diminished with knockdown of *TGFB1* (p = 0.0031), *TGFB2* (p = 0.0042) and *TGFB3* (p = 0.0183). Additionally, knockdown of *TGFBR2* (p = 0.0015) and *TGFBR3* (p = 0.0195) also significantly diminished production of pro-collagen I alpha, suggesting that the fibrogenic protein production in monoculture was downstream of Notch signaling-induced TGFβ. Subsequently, we examined how DLL4 stimulation regulated the distinct EC-proximal and EC-distal fibrogenic programs marked by *POSTN* and *COMP*, respectively (**Fig. 4k-l**). Although both genes were TGFβ responsive (p < 0.001), POSTN protein production was potentiated by Notch signaling (56-fold change, p = 0.003) whereas COMP protein production was suppressed (−2-fold change, p = 0.0082), implicating an additional mechanism by which Notch regulates the patterning of TGFβ responsiveness for the EC-distal gene *COMP*.

To quantify the dynamics of endothelial cell-derived Notch in regulating the EC-proximal and EC-distal fibrogenic programs, we first set up co-cultures with a fixed number of fibroblasts and varying numbers of endothelial cells; we collected and replaced media every 24 hours to measure the interval production of POSTN and COMP. POSTN protein production was detected at low levels in monocultured fibroblasts but was significantly induced in a EC ratio-dependent manner with an increasing number of ECs, starting as early as day 1 (p = 0.0089), through the duration of co-culture (p <0.0001) (**Fig. 5a**). In contrast, COMP protein production was significantly suppressed in EC:fibroblast co-cultures in a EC ratio-dependent manner compared to monocultured fibroblasts, starting as early as day 2 (p = 0.0432) and sustained throughout the time course (p = 0.0005) (**Fig. 5b**). Furthermore, the suppression of COMP production mirrored the EC:fibroblast ratio-dependent downregulation of TGFβRII (p < 0.0001, one-way ANOVA) and TGFβRIII (p < 0.0001, one-way ANOVA) suggesting that changes in fibroblast TGFβ receptor availability create dynamic threshold on the ability of fibroblast to respond to TGFβ (**Fig. 5c-d**).

**Figure 5.**
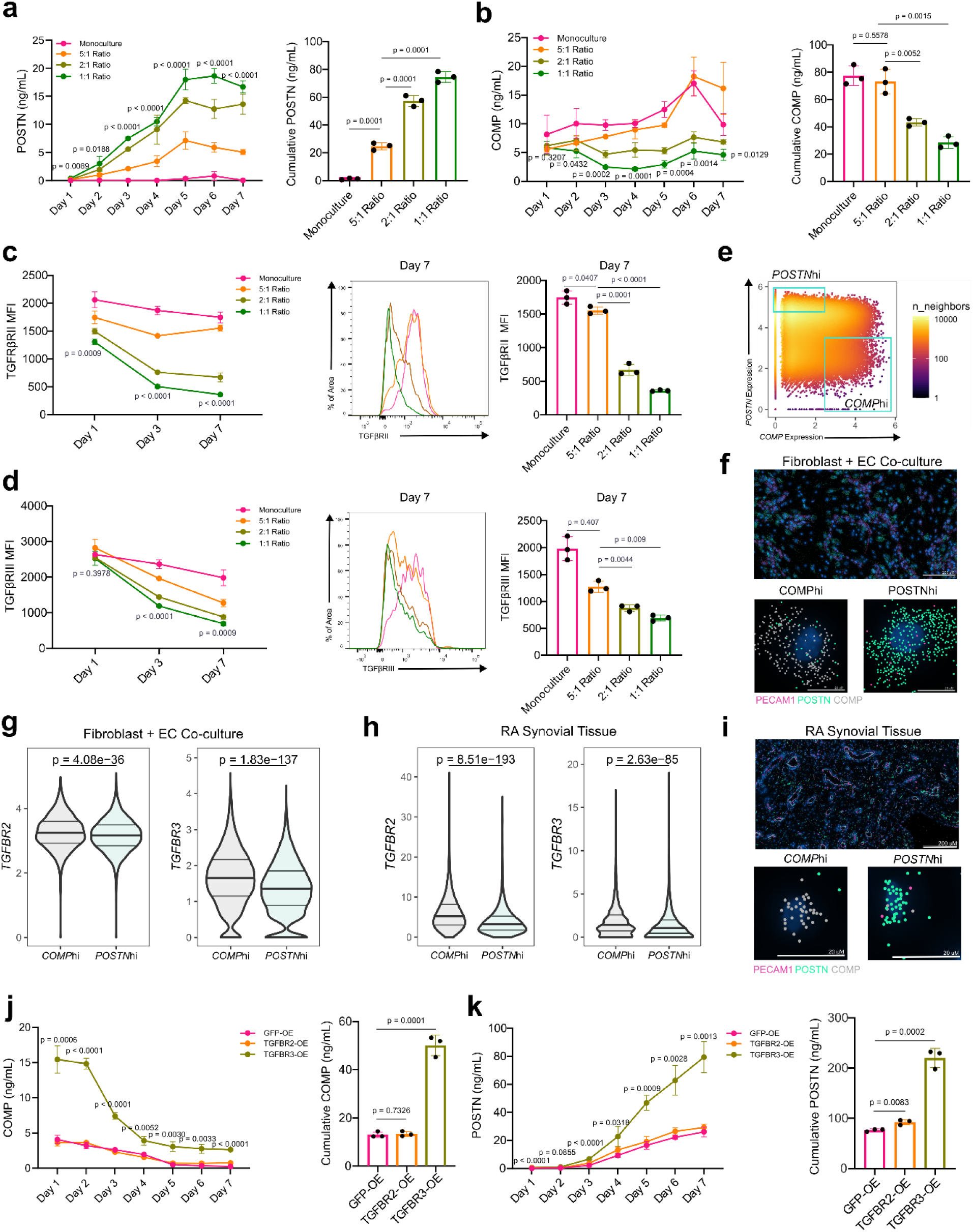
Endothelial-derived Notch signaling dictates fibroblast TGFβ responsiveness via regulation of TGFβ receptor III. **a-d,** A fixed number of fibroblasts were seeded with varying numbers of HUVECs. Monoculture: 5,000 fibroblasts, 5:1 ratio: 5,000 fibroblasts and 1,000 HUVECs, 2:1 ratio: 5,000 fibroblasts and 2,500 HUVECs, 1:1 ratio: 5,000 fibroblasts and 5,000 HUVECs. **a-b,** ELISA quantification of POSTN (a) and COMP (b) production over the course of co-culture. P-values in the line graph are displayed for the comparisons between monoculture and 1:1 ratio. The respective bar charts to the right represent the area under the curve. **c-d,** Flow cytometric quantification of fibroblast TGFβRII (c) and TGFβRIII (d) at the indicated days during co-culture with endothelial cells. P-values are shown for comparisons between monoculture and 1:1 ratio. Representative flow cytometry histograms and MFI quantification for day 7 of co-culture are shown. **e,** Gating strategy for classifying *COMP*hi and *POSTN*hi fibroblasts from the Xenium-profiled co-culture (mutually exclusive top quantile of cells expressing each gene, with *PO*STN ≤ 3.5 for *COMP*hi in this experiment). **f**, Representative examples of *COMP*hi and *POSTN*hi fibroblasts are shown with transcripts. **g,** Violin plot showing the distribution of *TGFBR2* and *TGFBR3* transcripts on *POSTN*hi and *COMP*hi gated fibroblasts in co-culture. **h**, Violin plot showing the distribution of *TGFBR2* and *TGFBR3* transcript expression in gated *POSTN*hi and *COMP*hi fibroblasts in synovial tissue. **i,** Representative example of gated (mutually exclusive top quantiles of expressing cells) *COMP*hi and *POSTN*hi synovial fibroblasts with transcr. **j-k**, ELISA quantification of COMP and POSTN production from co-culture of fibroblasts over-expressing (OE) GFP, TGFBR2 or TGFBR3 with endothelial cells in a 1:1 ratio. P-values are shown for the comparison between TGFBR3-OE and GFP-OE data points. The bar plot on the right represents the area under the curve. For **a-d,j-k,** Data points are shown as mean +/- s.d., represent n =3 biological replicates, and are representative of at least two independent experiments. For statistical analysis, two-tailed student’s t-test was used for **a-d, j-k** and two-sided Mann-Whitney U test was used for **f-i**.

To evaluate whether TGFβ receptor availability corresponded with COMP expression in fibroblasts, we further analyzed Xenium co-culture data and synovial tissue data, gating on *COMP*hi and *POSTN*hi cells and evaluated TGFβ receptor transcript expression in these populations (**Fig. 5e-i**). *COMP*hi fibroblasts had significantly higher levels of *TGFBR2* (co-culture: p = 4.06e-36, synovial tissue: p = 8.51e-193) and *TGFBR3* (co-culture: p = 8.51e-193, synovial tissue: p = 2.63e-85) compared to *POSTN*hi fibroblasts.

Next, to determine the extent to which suppression of COMP production in EC-fibroblast co-culture was mediated by dynamic TGFβ receptor downregulation, we used lentivirus to overexpress TGFBR2 and TGFBR3 in fibroblasts to restore TGFβ receptor expression (**Fig 5j-k, Extended Data Fig. 4a**). ECs co-cultured with fibroblasts overexpressing TGFBR3, but not TGFBR2, had significantly diminished suppression of COMP production, starting from day 1 (p = 0.0006), and maintained for the duration of co-culture (50.12 ng/mL vs. 13.02 ng/mL, 3.85-fold change, p = 0.0001). Further, daily POSTN production was significantly enhanced in co-cultures with TGFBR3-overexpressing, but not TGFBR2-overexpressing, fibroblasts starting day 1 (p <0.0001) and maintained throughout the course of co-culture (220 ng/mL vs. 76.39 ng/mL, fold change 2.88, p = 0.0002). Interestingly, overexpression of TGFBR2 and TGFBR3 had negligible effects on fibroblast response to TGFβ itself, suggesting that modulation of TGFBR3 availability is the specific mechanism by which endothelial-derived Notch signaling patterns TGFβ responsiveness (**Extended Data Fig. 4b**).

### Disruption of steady-state Notch patterning activates a fibrogenic program

Since we established that steady-state EC-derived Notch signaling generates a proximal to distal axis of TGFβ response in fibroblasts, we next sought to understand the consequences of disrupting steady-state patterning on the genesis fibrogenic gene expression. We hypothesized that disruption of a steady-state Notch signaling can activate expression of EC-distal gene expression programs within the EC-proximal compartment. We used siRNA to knock down EC-specific Notch ligand DLL4, fibroblast expressed JAG1 or a combination of both at the start of co-culture, collecting and replacing media daily for protein quantification (**Fig. 6a-b**). In all three conditions of Notch ligand knockdown, we observed significantly diminished production of POSTN by day 3 (siDLL4, p = 0.0126) compared to control, which was sustained for the duration of the time course (**Fig. 6a)**. COMP production was suppressed through day 3 for all conditions; however, starting on day 4, COMP production increased in the DLL4 knockdown condition (p = 0.0109) and stayed elevated compared to the control condition in which daily COMP production continued to decrease (p = 0.0081) (**Fig. 6b**). The effect of endothelial-specific DLL4 knockdown was much stronger than JAG1 knockdown, in which COMP production did not significantly increase, potentially due to active EC migration and proliferation throughout co-culture that results in a higher number of new endothelial-fibroblast contacts compared to fibroblast-fibroblast contacts.

**Figure 6.**
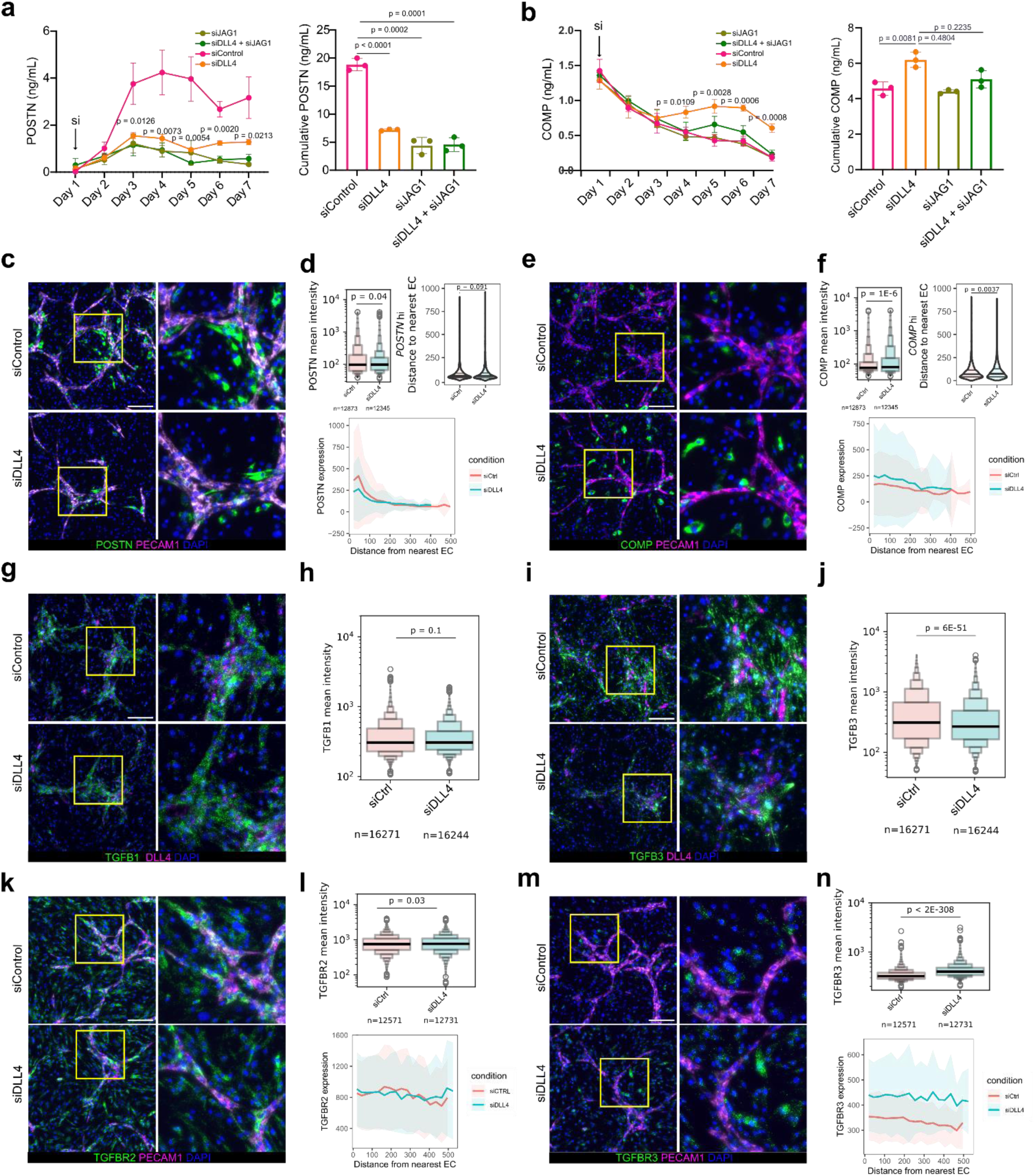
Disruption of steady-state Notch patterning activates a *COMP*hi fibrogenic program. **a-b,** ELISA quantification of POSTN (a) and COMP (b) production from fibroblasts and endothelial cell co-culture (1:1 ratio) treated with the indicated siRNAs (20 nM). P-values are shown for the comparison between DLL4 siRNA and control siRNA conditions. The bar plots to the right represent the area under the curve. **c-n**, RNAscope quantification of fibrogenic program in response to Notch perturbation. Representative images of siControl-treated and siDLL4-treated co-cultures are shown for the indicated genes with *PECAM1* and *DLL4* as endothelial markers; scale bar indicates 200 μm. **d,f,h,j,l,n**, Boxen plots represent the distribution of mean fluorescent intensities of the corresponding genes in siControl and siDLL4 conditions on the indicated number of cells selected for quantification. **d,f,l,n** Ribbon plots showing spatial gene expression patterns in relation to EC proximity for indicated genes between siRNA conditions. Solid line shows mean, shaded regions show one standard deviation. **d,f** Violin plots showing the distance to the nearest EC (in μm) for *POSTN*hi and *COMP*hi cells, defined as cells with the highest quantile of *POSTN* and *COMP* fluorescence intensity respectively. The data in **a-b** are shown as means ± s.d. and, represent n = 3 biological replicates and are representative of at least two independent experiments. A two-tailed unpaired student’s t-test was performed for **a**,**b** and two-tailed unpaired Mann-Whitney U tests for **d,f,h,g,l,n**.

Next, to characterize alterations in spatial patterning resulting from disruption of patterning generated by steady-state Notch signaling, we applied *in situ* hybridization to co-cultures treated with control siRNA or DLL4 siRNA (**Extended Data Fig. 5a)**. As captured by ELISA, we observed significant induction of *COMP* expression (p = 1.4e-6) and diminishment of *POSTN* expression (p = 0.04) in the DLL4 knockdown condition, quantified by comparing the mean gene expression intensities of fibroblasts in both conditions (**Fig 6c-f**). When we compared the average expression of *POSTN* in relation to EC distance, we observed no significant changes, as expected, given its EC-proximal restricted patterning (p = 0.0913) (**Fig. 6c-d**). In contrast, we observed that average *COMP* expression was higher compared to control in both the EC-proximal and EC-distal layers of the DLL4 knockdown condition, and that the average distance of a *COMP*hi cell was significantly higher in DLL4 knockdown (p = 0.00367) consistent with the broad spatial expansion of *COMP* expressing fibroblasts (**Fig. 6e-f**). We also examined the effect of DLL4 knockdown on TGFβ isoform and receptor expression (**Fig 6. g-n**). While we observed no significant changes in the expression patterns of *TGFB1* (p = 0.1) and only modest changes for *TGFBR2* (p = 0.03) expression patterns (**Fig 6. g-h, k-l**), we observed a highly significant increase in the average expression of *TGFBR3* (p < 2e-308) and decrease in the average expression of *TGFB3* (p = 6e-51) (**Fig 6. i,j,m,n**). As with *COMP* expression, we observed substantially elevated *TGFBR3* expression in both EC-proximal and EC-distal compartments of DLL4 knockdown co-culture (**Fig. 6n**). Together, these data suggest that the expansion of *COMP*-expressing fibroblasts was specifically linked to de-repression of TGFβRIII, mediated by DLL4 knockdown, rather than increases in TGFβ expression. Overall, this data supports the ability of EC-derived Notch signaling to dictate the spatial patterning of fibrogenic signaling by tuning fibroblast TGFβ sensitivity.

To examine the extent to which *COMP* expression was linked to Notch-mediated tuning of fibroblast sensitivity *in vivo*, we extended our analysis of RA synovial spatial data generated from the 396.10 cohort. We previously confirmed elevated *TGFBR2* and *TGFBR3* transcripts on *COMP*hi fibroblasts, consistent with the de-repression of *COMP* and *TGFBR2/3* at lower Notch-signaling levels (**Fig. 5j**). We then scored vascular niches according to their expression of Notch transcriptional targets (*HES1, HEY1, JAG1, NOTCH3, DLL4, NOTCH4, NOTCH1, COL4A1* and *COL4A2*). Across tissue samples, Notch activation score per vascular niche strongly negatively correlated with *TGFBR3* expression, although *COMP* expression showed no significant correlation with Notch activity possibly due to the heterogeneity of vascular subtypes associated with *COMP*hi fibroblasts (**Extended Data Fig. 5b**). Collectively, our *in vitro* and *in vivo* results suggest that Notch can tune TGFβ sensitivity via TGFβ receptor regulation in vascular niches.

### Spatiotemporal evolution of the *COMP*hi fibrogenic program

To characterize the spatiotemporal dynamics of fibroblast *COMP* expression in relation to treatment response, we examined changes in the abundance and localization of *COMP*hi fibroblasts in 17 paired post-treatment synovial biopsies. Across almost all synovia, including non-remission samples, we observed significant immune depletion post-treatment, regardless of the specific treatment received (**Fig. 7a-b, Extended Data Fig. 6a**). In contrast, we observed increases in the percent of fibroblasts expressing *COMP* in most patients, with starker increases under DMARD treatment than Adalimumab (**Fig. 7c-d**). Independent of treatment received, 5/6 non-remission patients showed an increase in *COMP* expression after 6 months, while half of remission patients (5/10) showed stable *COMP* expression (Δ*COMP*+ fibroblasts < 1%), suggesting that the expansion of the *COMP*hi fibroblast compartment is more frequent in non-responders. To better understand changes in the spatial characteristics of *COMPhi* tissue regions over time, we analyzed the cell densities of these regions pre- and post-treatment and observed that *COMPhi* regions tended to become more pauci-cellular post-treatment (**Fig. 7e-g**); the accumulation of *COMP*+ fibroblasts post-treatment negatively, but non-significantly, correlated with the cell density of *COMP*hi regions (R = −0.44, p = 0.073, two-sided Pearson correlation test) (**Fig. 7g**). Together, these data suggest that *COMP-* expressing fibroblasts expand post-treatment and accumulate in pauci-cellular niches over time, providing a first indication of active fibrogenic remodeling in the fibroblast-rich compartment of RA.

**Figure 7.**
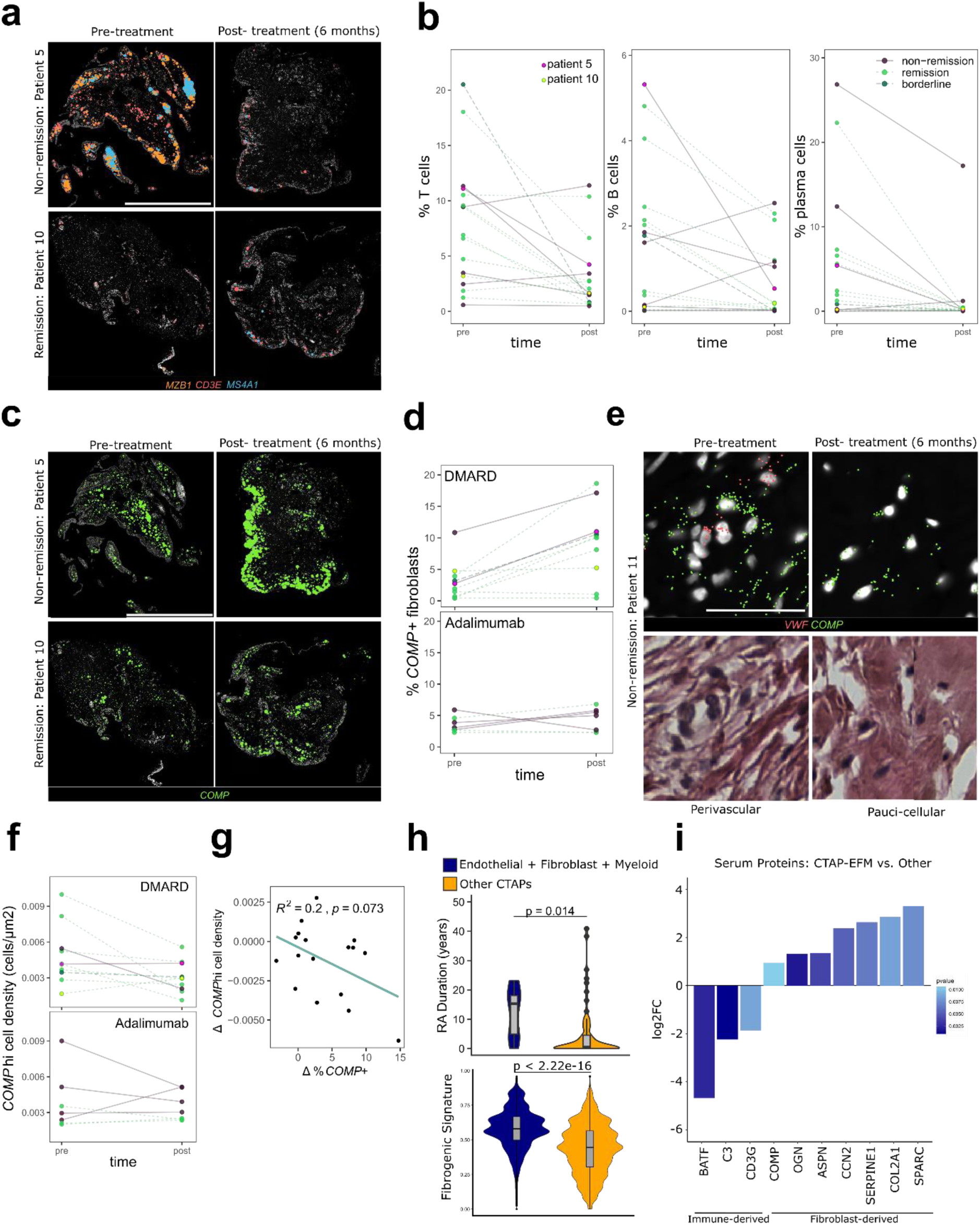
Spatiotemporal evolution of the *COMP*hi fibrogenic program pre- and post-treatment. **a,** Representative pre- and post-treatment gene expression from patients in remission (patient 10) or non-remission (patient 5) at 6 months. *MZB1* marks plasma cells, *CD3E* T cells and *MS4A1* B cells; scale bar indicates 2 mm. **b,** Quantification of the relative abundance of immune cell types pre- and post-treatment. Line colors indicate remission status. Patient 5 is highlighted in magenta, and Patient 10 is highlighted in yellow. **c,** Representative examples of *COMP* transcript expression in pre- and post-treatment synovial biopsies for remission and non-remission. **d,** Percent of fibroblasts expressing *COMP* per patient, pre- and post-treatment, separated by treatment type. **e,** Representative *COMP* gene expression and paired H&E for a pre- and post-treatment biopsy from a non-remission patient. *VWF* marks vasculature; scale bar indicates 50 uM. **f,** Cell density of *COMPhi* regions of the synovium (defined by kernel density estimate) per patient, pre- and post-treatment. **g,** Correlation between changes in the percent of fibroblasts that express *COMP* and the cell density of *COMP*hi regions per patient after 6 months of treatment. **h,** Violin plot representing the distribution of disease duration in RA patients classified as CTAP-EFM (n = 7) versus RA patients classified as other CTAPs (n = 63) and a violin plot representing the distribution of fibrogenic score, as calculated with UCell, of synovial fibroblasts from CTAP-EFM patients compared to fibroblasts from RA patients classified as other CTAPs. P-value was calculated with two-sided Mann Whitney U test for RA duration and a two-sided Wilcoxon test was used for comparison of fibrogenic score between groups. **i,** Bar plot of selected serum proteins that are differentially abundant in RA patients classified as CTAP-EFM compared to RA patients classified as other CTAPs. The height of the bar represents the log-fold change, and the shading of the bar represents unadjusted p-values.

Lastly, we evaluated whether serum markers could be used as surrogates to track fibrogenic remodeling in synovium. To do so, we further analyzed synovial fibroblast data^15^ to identify which cell-type abundance phenotype (CTAP) was most associated with fibrogenic signaling (**Fig. 7h-i**). Fibroblasts from the CTAP characterized by an abundance of endothelial, fibroblast and myeloid cells (CTAP-EFM) were the most strongly enriched for the fibrogenic gene signature and *COMP* expression compared to fibroblasts from other CTAPs (**Extended Data Fig. 6b**). Interestingly, CTAP-EFM patients, on average, also had significantly longer disease duration (p = 0.014) (**Fig. 7h**). To identify proteomic signatures associated with CTAP-EFM, we analyzed serum data, that was generated using a ∼5400-plex protein panel (Olink), from patients within the AMP cohort and compared the differential abundance of proteins in CTAP-EFM patients (n = 7) versus those classified as belonging to other CTAPs (n = 60). In CTAP E-F-M patients, we observed surprising enrichment of several established fibrogenic markers including *SPARC, OGN* and *CCN2*^36–38^ (**Fig. 7i, Extended Data Fig. 6c**). Collectively, the data supports fibrogenic remodeling in a subset of longer disease-duration RA patients and the potential to capture serum fibrogenic signatures systemically^39^.

## Discussion

Treatment failures in RA lead to high costs to society and cause undesirable morbidity associated with immunosuppression. Several factors have been suggested to contribute to RA treatment failure^3^, yet no novel therapeutically targetable mechanism has been identified for refractory RA. Our study defined a Notch-TGFβ-driven fibrogenic transcriptional program associated with non-remission in RA. Our result is consistent with previous studies that have suggested an association between the expansion of synovial fibroblasts and refractory RA^15,17,40^. A recent study uncovered a reciprocal relationship between perivascular fibrogenic fibroblasts and immune cell invasion^41^. Interestingly, though, we find that lower immune cell invasion does not always lead to the resolution of disease. Higher *COMP* expression in treatment-naïve biopsies of non-remission patients suggests that the baseline state of synovial tissue contributes to fibrogenic tissue remodeling. By implicating Notch signaling in regulating fibroblast TGFβ activity, our results provide a molecular mechanism for synovial tissue fibrosis in RA.

Notch is an evolutionary-conserved pathway by which cells generate borders and boundaries^42^. In fibroblasts, we previously reported an essential role of Notch signaling in the determination of vascular fibroblast positional identity^9^. Our results demonstrate the tight control of TGFβ receptor signaling in the perivascular compartment downstream of Notch. Our model further suggests that the border between mural cells and fibroblasts is actively maintained through steady-state Notch signaling from the vascular endothelium and that disruption or loss of steady-state Notch signaling resensitizes perivascular fibroblasts to TGFβ signaling, generating the fibrogenic niche. In this model, tissue fibrosis occurs when the system senses a loss of vascularization through the loss of Notch signaling.

Consistent with previous studies of RA synovia using single-cell approaches^15^, we find that immune cells particularly decrease in abundance with treatment. However, we find this occurs whether patients achieve remission or not, suggesting that reducing the immune compartment alone is not sufficient to reliably resolve RA. Several recent studies have highlighted pain-sensing as an important feature of synovial biology, and that a subset of RA patients experience persistent pain despite minimal evidence of synovial inflammation^43,44^. We hypothesize that fibrogenic remodeling in the synovium may contribute to joint stiffness and pain as has been described in OA^19,39^. Based on our findings, targeting fibrogenic activation and endothelial-fibroblast crosstalk may interrupt synovial tissue fibrosis and prevent refractory RA. Specifically, our study implicates TGFβ3 and TGFβRIII as spatially regulated components of TGFβ signaling that drive differential TGFβ response in fibroblasts. Of note, therapeutic targeting of TGFβ3 by anti-TGFβ3 antibody attenuated fibrosis in an animal model of fibrosis^45^, raising the possibility of selectively targeting TGFβ3 to prevent exuberant synovial tissue fibrosis in RA patients.

In this cohort we also had the ability to assess differential effects of first line triple DMARD therapy versus TNFi in recent-onset RA, which is not typical standard of care. Triple DMARD therapy appeared to be as or more effective than the anti-TNF biologic in depleting immune populations; however, we observed greater expansion of the treatment-resistance associated *COMP+* fibrogenic compartment in patients treated with DMARDs. We hypothesize that the sub-population of patients who achieved remission at 6 months but still showed increased *COMP* expression may have a higher risk of relapse compared to patients with stable levels of *COMP*, which is consistent with our finding that the *COMP*+ fibroblast population co-expresses genes previously reported to be enriched in RA patients refractory to multiple lines of therapy^17^. We also observe a trend towards lower cell density within *COMP*hi regions over time, which could be the first indication of a progression from the vasculogenic initiation of fibrosis to pauci-cellular fibrotic bands that continue to express *COMP* and other ECM components. A validation study with a larger cohort and longitudinal data by the Accelerating Medicines Partnership Autoimmune and Immune-Mediated Disease network is underway to address this question.

## Methods

### Human clinical samples

RA synovial tissue was obtained from patient donors at the Flinders Medical Center (Protocol #396.10), Brigham and Women’s Hospital (IRB no. 2019P002924) and University of Colorado Hospital (IRB 20-1908.) Samples were obtained from synovial biopsies or retrieved from RA patients undergoing arthroplasty or synovectomy and were FFPE or OCT-embedded immediately after retrieval. Clinical characteristics of patients are available in Supplementary Table 1. Enrollment complied with relevant ethical regulations of all institutions. Informed consent was obtained from all participants.

### Xenium slide preparation

FFPE blocks were sliced onto Xenium slides according to the “Xenium In Situ for FFPE-Tissue Preparation Guide” (CG000578 Rev C, 10X Genomics) protocol. The Xenium slides were then prepared following the “Xenium In Situ for FFPE-Deparaffinization and Decrosslinking” protocol (CG000580 Rev D, 10X Genomics). In brief, the slides were baked at 60°C for 30min and then sequentially immersed in xylene, ethanol, and nuclease-free water to deparaffinize and rehydrate the tissue. Immediately after, the Xenium slides were incubated in the decrosslinking and permeabilization solution at 80°C for 30 minutes, followed by a wash with PBS-T.

For co-cultures, Xenium slides were pre-coated with diluted1:10 diluted matrigel (Corning, cat. 356231) after which a mixture of fibroblasts (120,000) and HUVECs (60,000) were seeded for 72 hours. Slides were fixed and permeabilized according to protocols for fresh frozen tissue (CG000581).

Next, the slides were prepared according to the “Xenium Prime In Situ Gene Expression” user guide (CG000760 Rev A, 10X Genomics) for the remaining steps or “Xenium In Situ Gene Expression - Probe Hybridization, Ligation & Amplification” (CG000582) for co-culture. Probes from the Xenium Prime 5K Human Pan Tissue & Pathways Panel (PN- 1000671, 10X Genomics) and the custom addon panels (S. table X) were hybridized to the samples at 50°C for 18 hours. Post-hybridization, the slides underwent washing, incubation with a ligation reaction mix, another wash step, and DNA amplification. Finally, cell segmentation staining was conducted, and the slides were treated with an autofluorescence quencher and DAPI before being loaded into the Xenium instrument.

### Xenium Analyzer setup and data acquisition

Processed Xenium slides were loaded in the Xenium Analyzer and imaged, following the guidelines in the “Xenium Analyzer User Guide (CG000584 Rev F, 10X Genomics).” After scanning, the Xenium slides were removed from the Xenium Analyzer, and processed with post-run H&E staining according to the “Xenium In Situ Gene Expression - Post-Xenium Analyzer H&E Staining” protocol (CG000613 Rev A, 10X Genomics).

### Xenium data cell typing and niche analysis

Cells in Xenium data were segmented using Xenium Onboard Analysis, based on nuclear expansion (custom panel co-culture data) or multimodal cell segmentation using cell boundary, interior, and nuclear stains (5k data).

Single-cell and niche analysis were performed using Seurat v5.0.0 after filtering out cells with ≤ 50 transcript species (“nFeatures”) and log normalizing data for each sample with the median transcripts detected as the scale factor. Single cells from different samples were integrated using Harmony^46^. Niches were identified using Tessera^23^ based on Harmony embeddings. We labelled cell types and niches in 5,101-gene panel data based on the enrichment of marker genes associated with co-clustering cells or niches. For cell types, neighbor detection and UMAP were run with 20 principal components, and clustering with a resolution of 0.3. For niches, neighbor detection and UMAP were run with 10 principal components, and clustering with a resolution of 0.2. Example niches were selected among the top three most typical (by Euclidean distance to the average cell type representation) within a single sample in which all niches were detected. In some cases, Xenium Explorer v3.2 was used for visualization of transcripts.

For the 50-gene custom panel data, endothelial and mural cells were identified based on enrichment of markers (*PECAM1* and *ACTA2*, respectively) in clustered data. For comparison to *COMP*+ fibroblasts, fibroblasts were identified based on expression of *PDGFRB* and *NOTCH2* > 0 and exclusion from endothelial and mural cell clusters.

### Transcript kernel density estimate analysis

We calculated kernel density estimates using the tidy_kde function from the eks R package ^47^. Pairwise Pearson correlations were calculated by taking density values for different transcripts at each pixel within a sample, after selecting for subregions on the slide based on 50% contour regions of the *COL1A1* kernel density estimate or regions with DAPI above background, as specified. For *COMP*hi regions in Figure 7, we considered cell counts per area within regions defined by the 50% contour region of the kernel density estimate per sample. We used the R packages terra and raster to mask subregions.

### scRNA-seq analysis

For analysis of scRNAseq data, we used the R package Seurat v5.0 for downstream analysis. The filtered count matrix was used as input, and we used SCTransform to normalize data prior to principal coordinate analysis and UMAP projection. Differential expression analysis was performed using the FindMarkers function with a Wilcoxon test and gene signature scores were applied to datasets using the AddModuleScore_Ucell function from the UCell package. The DotPlot function was used to generate bubble plots depicting the expression of indicated genes across cell populations and the VlnPlot function was used to generate violin plots.

### Immunofluorescence

OCT embedded synovial sections were equilibrated at −20C for 10 minutes, and 10-mm sections were obtained using a cryostat, mounted onto a glass slide and stored at −80C. Prior to staining, slides were equilibrated for 5 minutes at room temperature, sections were encircled with a PAP pen (ImmEdge, Vector Labs), rehydrated with PBS for 5 minutes at RT, fixed for 10 minutes in 4% PFA, washed three times with PBS, and blocked for 1 hour in TBS-T with 5% BSA and 0.2% Triton X-100. The following primary antibodies were diluted in blocking buffer and incubated overnight at 4°C: phospho-SMAD3 (Thermofisher, 1:200 dilution), TGF beta-1 (Thermofisher 1:100), TGFB beta-2 (Thermofisher, 1:50), TGF beta-3 (Thermofisher, 1:100), NOTCH3-PE (Biolgend, 1:50), CD45-FITC (Biolegend, 1:50), Collagen I (Thermofisher, 1:1000) and VWF (Thermofisher, 1:100). Primary antibodies were removed, and sections were washed with PBS-T three times prior to secondary antibody incubation with the following at a 1:500 dilution in blocking buffer: AF660 anti-mouse (Thermofisher, A-21074) and AF555 anti-rabbit (Thermofisher, A-21424). Secondary antibodies were removed, sections were washed three times with PBS-T and mounted using Fluoromount-G mounting medium with DAPI (Thermofisher). Images were acquired on an EVOS M7000.

### In vitro culture

Synovial fibroblast cell lines were thawed and cultured in DMEM supplemented with 10% FBS, 10 mM HEPES, 1% MEM amino acids, 1% non-essential amino acids, 1% L-glutamine, 1% penicillin/streptomycin, 55 uM β-mercaptoethanol and 1% gentamicin. HUVECs (Thermofisher) were thawed and maintained in EGM-Plus media supplemented with the EGM-plus bulletkit (Lonza). Passages 3 to 6 were used for experiments. For DLL4 stimulation experiments, cell culture plates were pre-coated overnight with 5 μg ml−1 DLL4-Fc (R&D Systems) and for co-cultures, plates were pre-coated overnight with a 1:10 dilution of matrigel (Corning, Cat. 356231) and washed with PBS. In some conditions, cells were treated with TGF beta-1 (Peprotech, 10 ng ml−1), DAPT (Selleckchem, 10 uM) or TGFβ inhibitor (SB431542). The following siRNAs (Thermofisher) were added to monocultures or co-cultures at a 20 nM concentration at the start of cultures and were re-dosed every 72 hours, when applicable: NTC (4390843), JAG1 (s1174) and DLL4 (s534448), TGFB1 (s14054), TGFB2 (s14059), TGFB3 (s224725), TGFBR1 (s14073), TGFBR2 (s14077), TGFBR3 (s24). siRNA treatment was performed as described in the Lipofectamine RNAiMax protocol (Thermofisher). For RT-qPCR experiments, 10,000 fibroblasts were seeded in 96-wells and for western blot 250,000 fibroblasts were seeded in 6 well plates prior to stimulation and cell lysis. For co-cultures a fixed number of fibroblasts (5,000) were cultured with HUVECs (5,000) at a 1:1 ratio, unless otherwise indicated in the figure legends.

### Flow cytometry

Fibroblasts or co-cultures were trypsinized and directly stained with an equal volume of primary antibody solution (Cell staining buffer, Biolegend) containing eFluor 780 Fixable Viability Dye (Thermofisher, 1:1000), TGFBR2-APC (Biolegend, 1:100), TGFBR3 (Proteintech, 20000-1-AP, 1:100) or CD31-FITC (Biolegend, 1:100) for 30 minutes at 4C. Cells were washed with cell staining buffer (Biolegend), fixed in 4% PFA for 10 minutes at RT, and stained with AF555 anti-rabbit (Thermofisher) for 30 minutes at 4°C for detection of TGFBR3. Cells were washed and resuspended in cell staining buffer. Data was acquired on a BD FACSCanto II and further analyzed with Flowjo version 10.

### RNAScope and quantification

A 3:1 ratio of fibroblasts and endothelial cells (7500 fibroblasts, 2500 HUVECs) was seeded on matrigel (Corning, 1:10 dilution) pre-coated 96-well plates (Agilent, cat. 204626-100). After culture and treatment with siRNA, cells were washed with PBS, fixed in 10% NBF and processed for staining and imaging according to the manufacturer’s protocol with the RNAScope multiplex fluorescent V2 assay (ACD Bio, SOP 45-009A). Images were acquired on an EVOS M7000.

Results were quantified by segmenting nuclei based on DAPI staining using Cellpose^48^ and performing nuclear expansion to approximate cell boundaries. We then used scikit-image^49^ to calculate mean intensities per cell.

### Western Blot

Following incubation, cells were trypsinized and rinsed with cold PBS. Cells were pelleted by centrifugation at 1500 rpm for 10 minutes. Cell pellets were resuspended in 100 µL of RIPA buffer (Thermo Fisher Scientific, #89900) supplemented with 0.1% Triton x-100, protease and phosphatase inhibitor mini-tablets (Thermo Fisher Scientific, #A32961) and a phosphatase inhibitor cocktail (Active Motif, #37492). The cell lysate was briefly vortexed at high speed to dissociate aggregates, incubated on ice for 30 minutes, and then centrifuged at high speed for 20 minutes at 4°C. Protein concentration was determined using the Pierce BCA Protein Assay Kit (Thermo Fisher Scientific, #23227). Between 30-50 µg of protein was subjected to SDS-PAGE (Bio-Rad, #1610734) and transferred to a PVDF membrane (Bio-Rad, #1704273) using the Bio-Rad TRANS-BLOT TURBO transfer system (Bio-Rad, #690BR324) with a dry transfer protocol for 30 minutes according to the manufacturer’s instructions.

Membranes were blocked for 15 minutes in Everyblot blocking buffer (Bio-Rad # 12010020) then incubated overnight at 4°C with primary antibodies against TGFβ1 (Proteintech, # 21898-1-AP, 1:500), TGFβ2 (Proteintech, # 19999-1-AP, 1:500 dilution), TGFβ3 (Proteintech, # 18942-1-AP, 1:500 dilution), TGFβR1 (RD Biosciences #AF3026, 1:300 dilution), TGFβR2 (Bioss #bs-0117R, 1:500 dilution), TGFβR3 (Cell Signaling Technology #2519S, 1:500), GAPDH (Thermo Fisher Scientific, #MA5-15738), or beta-actin (Cell Signaling Technology, #3700). Following primary antibody incubation, membranes were incubated with HRP-conjugated secondary antibodies (Thermo Fisher Scientific, anti-Rabbit #32460, anti-Mouse #31430, or anti-Goat#A16005) for 1 hour at room temperature. All the antibody dilutions were in Everyblot blocking buffer. Blots were developed using SuperSignal™ West Femto Maximum Sensitivity Substrate (Thermo Fisher Scientific, #34095) or SuperSignal™ West Pico PLUS Chemiluminescent Substrate (Thermo Fisher Scientific, #34577) and imaged using a Bio-Rad ChemiDoc imaging system.

### Analysis of Olink Data

Olink protein concentration values (NPX) from AMP RA/SLE patients were extracted with the OlinkAnalyze package. A count matrix and a separate file with metadata indicating eaach patient’s CTAP was formed. The lmfit and eBayes functions from the limma package were used to identify the differential abundance of proteins in CTAP-EFM patients compared to patients classified in other CTAPs. Unadjusted p-values are reported.

### Bulk RNA-sequencing analysis

We analyzed both bulk RNA-seq and pseudo-bulk from spatial transcriptomics data using DESeq2^50^.

### ELISA

Supernatant was collected on the indicated days and stored at 4**°**C in sealed polypropylene plates prior to assay by ELISA. For measurement of Pro-Collagen 1 alpha 1 (R&D Systems, DY6220-05), COMP (R&D Systems, DY3134), or POSTN (R&D Systems, DY3548B) plates were incubated overnight at 4**°**C with diluted capture antibody solution, washed with PBS-T and blocked for 1 hour at RT in blocking buffer (1% BSA in PBS). Supernatant was diluted in blocking buffer and incubated on plates for 2 hours at room temperature, followed by a 2-hour incubation of detection antibody and 30-minute incubation of HRP-Streptavidin (R&D Systems) with PBS-T washes in between. The ELISA was developed with 50 µL of TMB substrate (Thermofisher) and stopped after 20 minutes with 50 µL of 2N sulfuric acid. Absorbance was read with a plate reader (Spectramax) at 450 nm with a 570 nm background subtraction.

### Quantitative polymerase chain reaction

Cells were lysed in TRIzol (Thermofisher) and RNA was extracted per manufacturer protocols. cDNA was synthesized using QuantiTect Reverse Transcription Kit (Qiagen) per manufacturer protocols. The qPCR mastermix was prepared using Brilliant III Ultra-Fast SYBR Green QRT-PCR Master Mix (Agilent) and samples were run on an AriaMX Real Time PCR machine (Agilent). The relative abundance of transcripts was normalized to the expression of glyceraldehyde-3-phosphate dehydrogenase (GAPDH) mRNA using the 2-ΔΔCT method.

### Lentiviral transduction

TGFBR2 and TGFBR3 plasmids (pLV[Exp]-Bsd-CMV) were ordered from VectorBuilder. 293FT cells were seeded at 6e5 cells in a T75 flask and incubated overnight. The next day, cells were transfected according the manufacturer’s protocol with 11.18 μg of target plasmid and 17.8 μg of packaging mix (Thermofisher, V53406) using Lipofectamine 3000 (Thermofisher, L3000015) in a T-75 flask. Virus supernatants were harvested and filtered after 4 days and concentrated overnight using Peg-it virus precipitation solution (System Biosciences) per manufacturer protocols. Cells were transduced with lentiviral supernatants containing polybrene (10 μg ml−1) for one week, with regular media changes, followed by a one week of selection with blasticidin (10 μg ml−1) prior to use in experiments.

### Statistics

Statistical tests were performed using Graphpad Prism version 10.4.1 or R version 4.4.1/4.4.2. There were no tests performed to assess normality. Differences between data that were assumed to be normally distributed were assessed using Student’s two-tailed t-tests and difference in data assumed to be non-normally distributed data were evaluated with Mann–Whitney U-tests. Detailed statistical descriptions, including sample size and calculation methods, are provided in the figure legends and text. A p-value of <0.05 was considered statistically significant.

## Acknowledgments

We thank members of the Wei lab, Korsunsky lab, and Brenner lab for helpful discussions. We also appreciate the contribution of synovial samples from the Colorado Interdisciplinary Joint Biology Program (CUIJBP) consortium to this study. This work is supported by a Doris Duke Foundation Clinical Scientist Development Award, a Rheumatology Research Foundation Innovative Research Grant, a Brigham and Women’s Hospital Department of Medicine - Broad Institution collaborative research Award, a Brigham and Women’s Hospital Llura Gund Award for Rheumatoid Arthritis Research. This work was supported by the Accelerating Medicines Partnership Autoimmune and Immune-Mediated Diseases Network (AMP-AIM). AMP-AIM is a public-private partnership (AbbVie Inc., Arthritis Foundation, Bristol-Myers Squibb Company, Foundation for the National Institutes of Health, GlaxoSmithKline, Janssen Research and Development, LLC, Lupus Foundation of America, Lupus Research Alliance, Merck Sharp & Dohme Corp., National Eye Institute, National Institute of Allergy and Infectious Diseases, National Institute of Arthritis and Musculoskeletal and Skin Diseases, National Institute of Dental and Craniofacial Research, National Institute Health Office of Research on Women’s Health, Norvartis, Pfizer Inc., Sanofi, and Sjogren’s Foundation, and UCB) created to develop new ways of identifying and validating promising biological targets for diagnostics and drug development Funding was provided through grants from the National Institutes of Health. K.W. is supported by a NIH-NIAMS K08AR077037, a Burroughs Wellcome Fund Career Awards for Medical Scientists. K.B. is supported by a NIH-NIAMS T32AR007530-36.

## Author contributions

Conceptualization: K.B., K.W., M.D.W., R.T. Experimental design and data generation: K.B, S.K., V.K. Xenium data generation and panel design: G.C., S.A.P, K.B., R.M., V.K. Data analysis: A.B.R.M., K.B., M.T., R.L, K.S.A. Synovial tissue acquisition: S.P., P.E.B., J.K.L., A.H.J, M.D.W., R.T. Analysis of published data: R.L., A.B.R.M., K.B., M.J.L., C.P. Figure generation: K.B., A.B.R.M. writing original draft: K.B., A.B.R.M., K.W. draft reviewing and editing: all. Supervision: K.W., M.D.W. Funding acquisition: K.W.

---

**Extended Data Figure 1.**
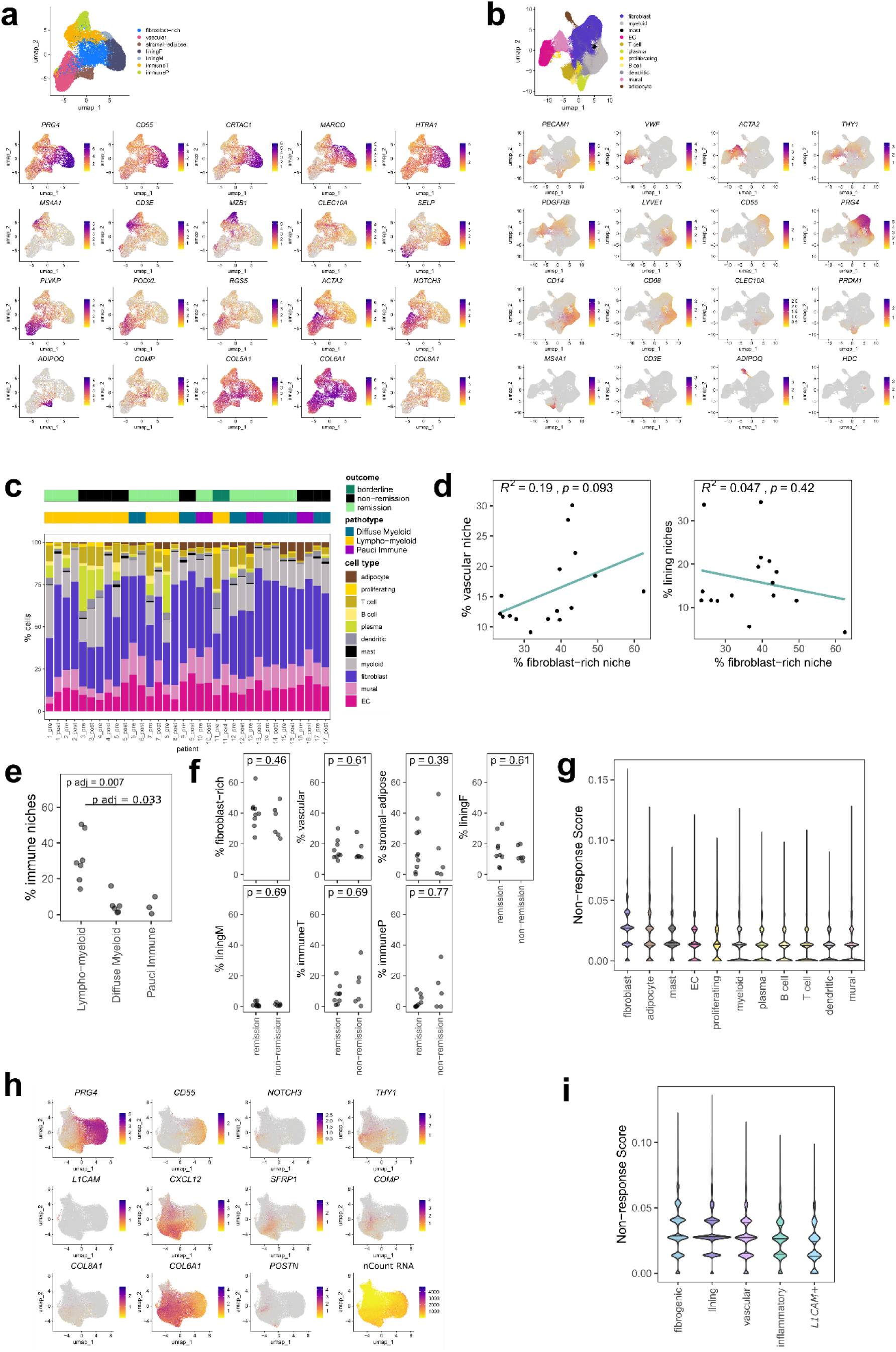
Cell and niche analysis of pre-treatment RA synovium. **a,** UMAPs showing labelled niches (above) and niche marker genes (below). Each point represents a single niche. **b,** UMAPs showing labelled cell types (above) and cell type marker genes (below). Each point represents a single cell. **c,** Stacked bar plot summarizing cell type proportion in pre- and post-treatment samples for 17 patients. **d,** Correlations between the percent area of each sample occupied by the fibroblast-rich versus vascular or lining niches. **e,** The percent area of each sample occupied by immuneT and immuneP niches by pathotype. **f,** Dot plot representing the tissue area of niche in baseline biopsies by remission status. **g,** Violin plot representing the distribution of non-response scores per cell type. **h,** UMAP displaying markers for different fibroblast subtypes labelled in Figure 1h. **i,** Violin plot representing the distribution of non-response scores per fibroblast subtype.

**Extended Data Figure 2.**
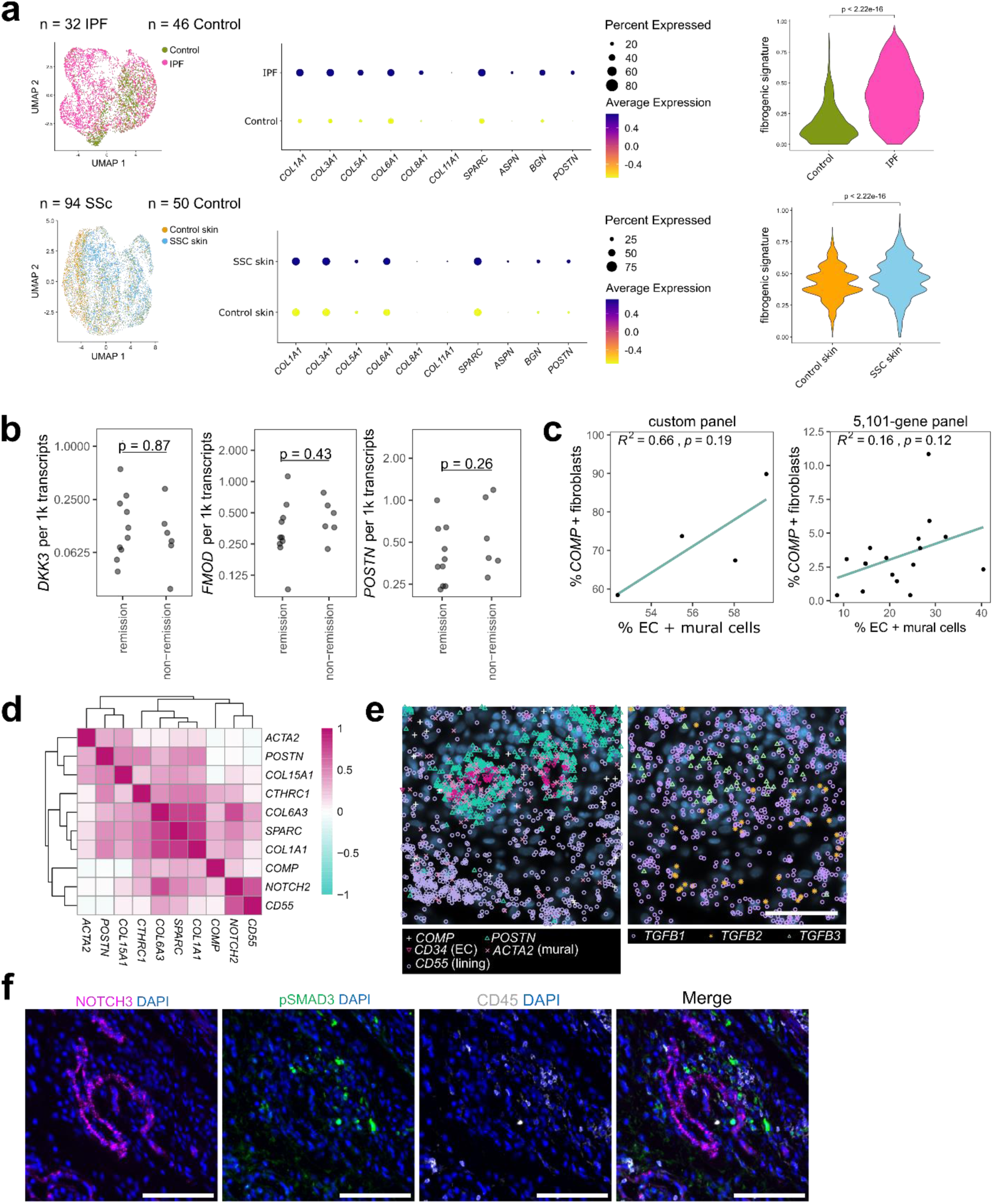
Analysis of fibrogenic program in RA synovium. **a,** UMAP projection of single cells from IPF and SSc datasets grouped by disease status (refs.^26,27^) and a corresponding dotplot of genes (n = 10) that comprise the fibrogenic gene signature. Dot plot color represents the scaled expression of genes, and the size of the dots represent the percent of cells expressing the gene. Violin plot displays the fibrogenic signature score, as calculated with UCell, in disease versus control samples. Wilcoxon test was used for statistical comparisons between the groups. **b,** Dot plot representing normalized transcript detection of *DKK3*, *FMOD*, and *POSTN* from pre-treatment spatial transcriptomics data by response status. **c,** Correlation between the percent of fibroblasts that express *COMP* (count > 0) per sample and abundance of endothelial and mural cells as a percent of total cells in the same sample. Left: 4 samples analyzed with a 50-gene custom panel. Right: 16 samples analyzed with a 5,101-gene panel. **d,** Heatmap representing spatial density correlations between fibrogenic and cell type marker genes (mean across 4 samples in the 50-gene custom panel). **e,** Examples of *TGFB* ligand transcript expression patterns in 50-gene custom panel, with cell type and subtype marker genes shown on the left for the same site. Scale bar indicates 50 µm. **f,** Representative immunofluorescence images showing pSMAD3 staining in CD45-cells in a NOTCH3+ perivascular region. Scale bars indicate 200 µm.

**Extended Data Figure 3.**
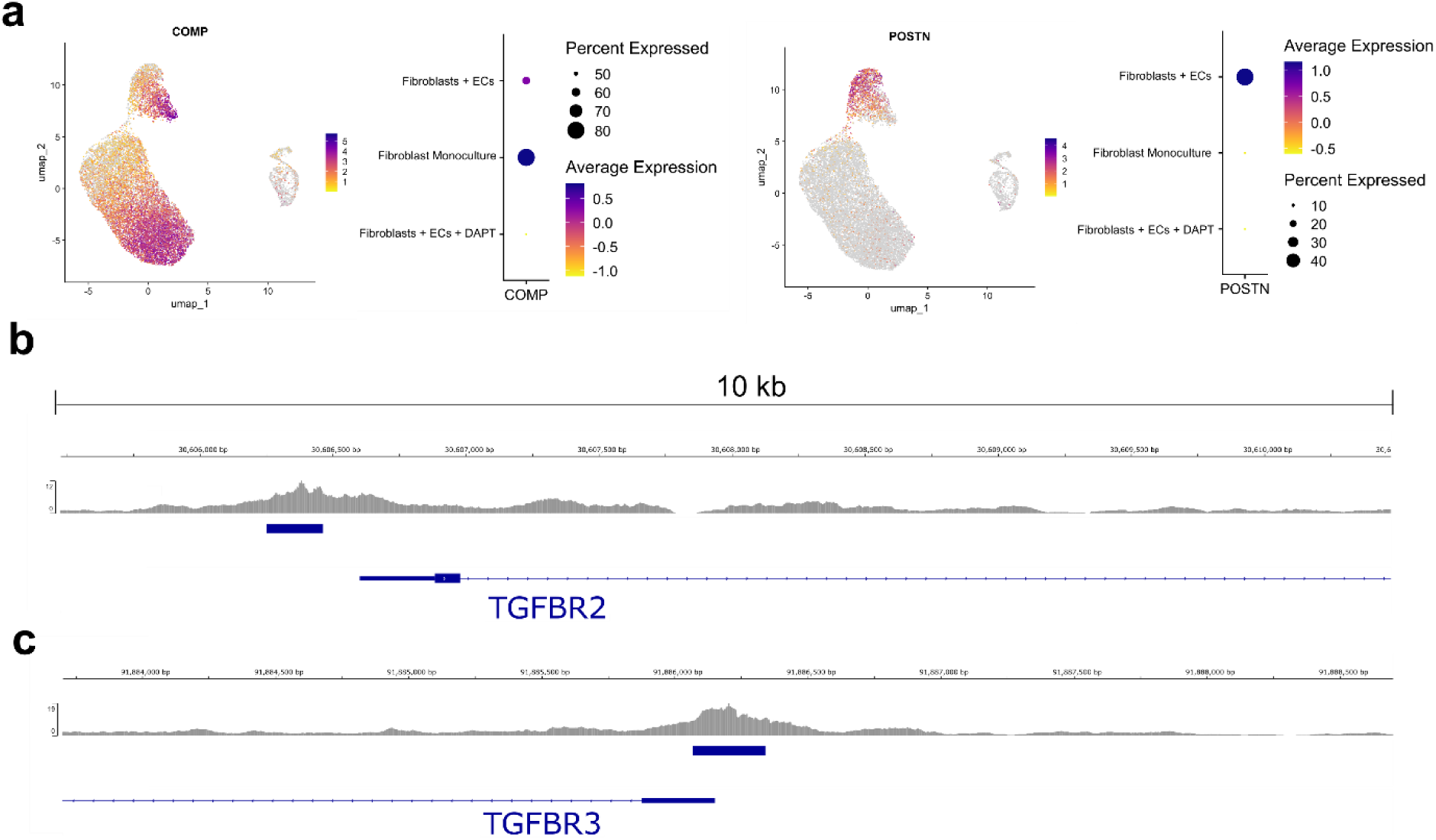
Micromass and ChIP-Seq analysis. **a,** UMAP projection and dotplots of micromass organoid data representing *COMP* and *POSTN* expression**. b-c**, Chip-seq analysis of RBPJ binding to *TGFBR2* and *TGFBR3* promoter regions in HepG2 cells (data from ENCSR596FEL). Significant peaks are annotated with blue bars and were called using the narrowPeak function in MACS2.

**Extended data Figure 4.**
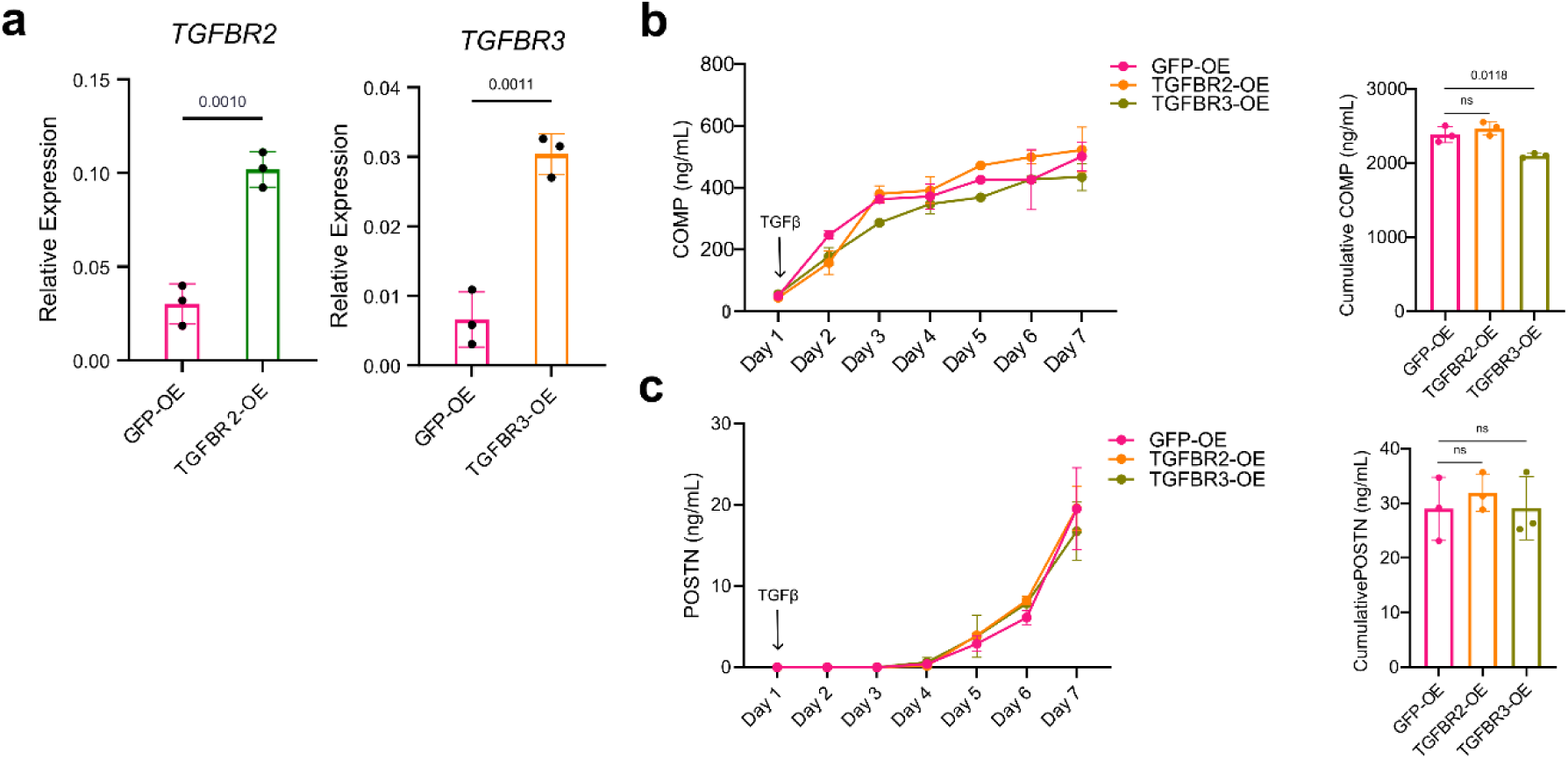
Lentiviral overexpression of TGFBR2 and TGFBR3. **a**, RT-qPCR analysis of *TGFBR2* and *TGFBR3* expression in TGFBR2-overexpressing or TGFBR3*-*overexpressing fibroblasts respectively, compared to control GFP-overexpressing fibroblasts. **b-c,** ELISA quantification of COMP and POSTN production from GFP-OE, TGFBR2-OE or TGFBR3-OE stimulated continuously with recombinant TGFβ1 (10 ng ml−1) stimulation. The bar plots to the right represent the area under the curve. For all comparisons, two tailed student’s t-test was used, data represents n = 3 biological replicates, are shown as mean +/- s.d. and data are representative of at least two independent experiments.

**Extended Data Figure 5.**
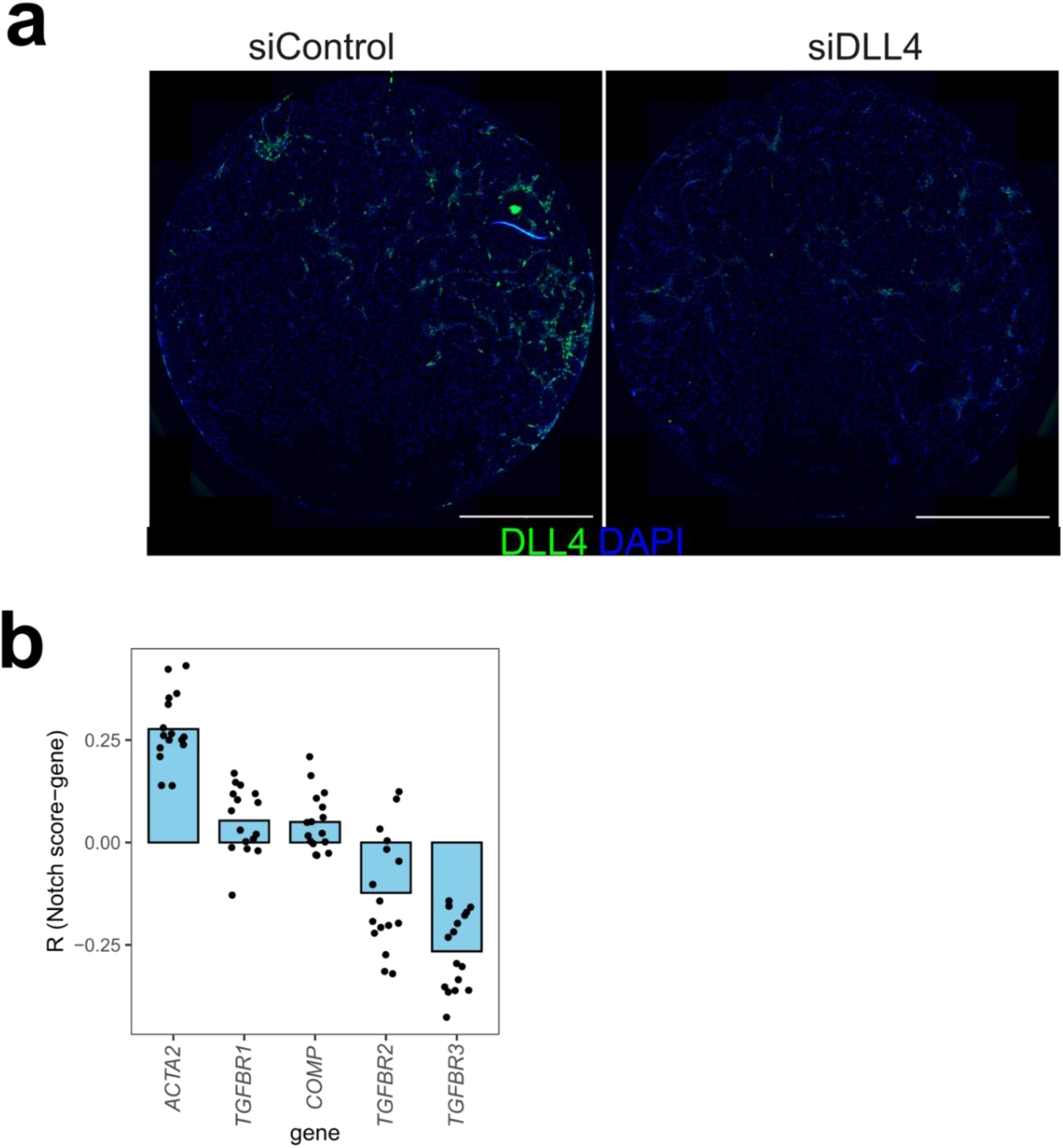
Validation of DLL4 knockdown and correlation of Notch activation score with vascular niche transcripts. **a,** Representative RNAscope image showing DLL4 expression in siControl and siDLL4-treated co-culture. Scale bar indicates 2mm. **b,** Mean Pearson correlation (R) between Notch activation score and transcripts detected per vascular niche for each pre-treatment sample(n = 16). *ACTA2* represents a positive control for expected correlation with Notch signaling.

**Extended Data Figure 6.**
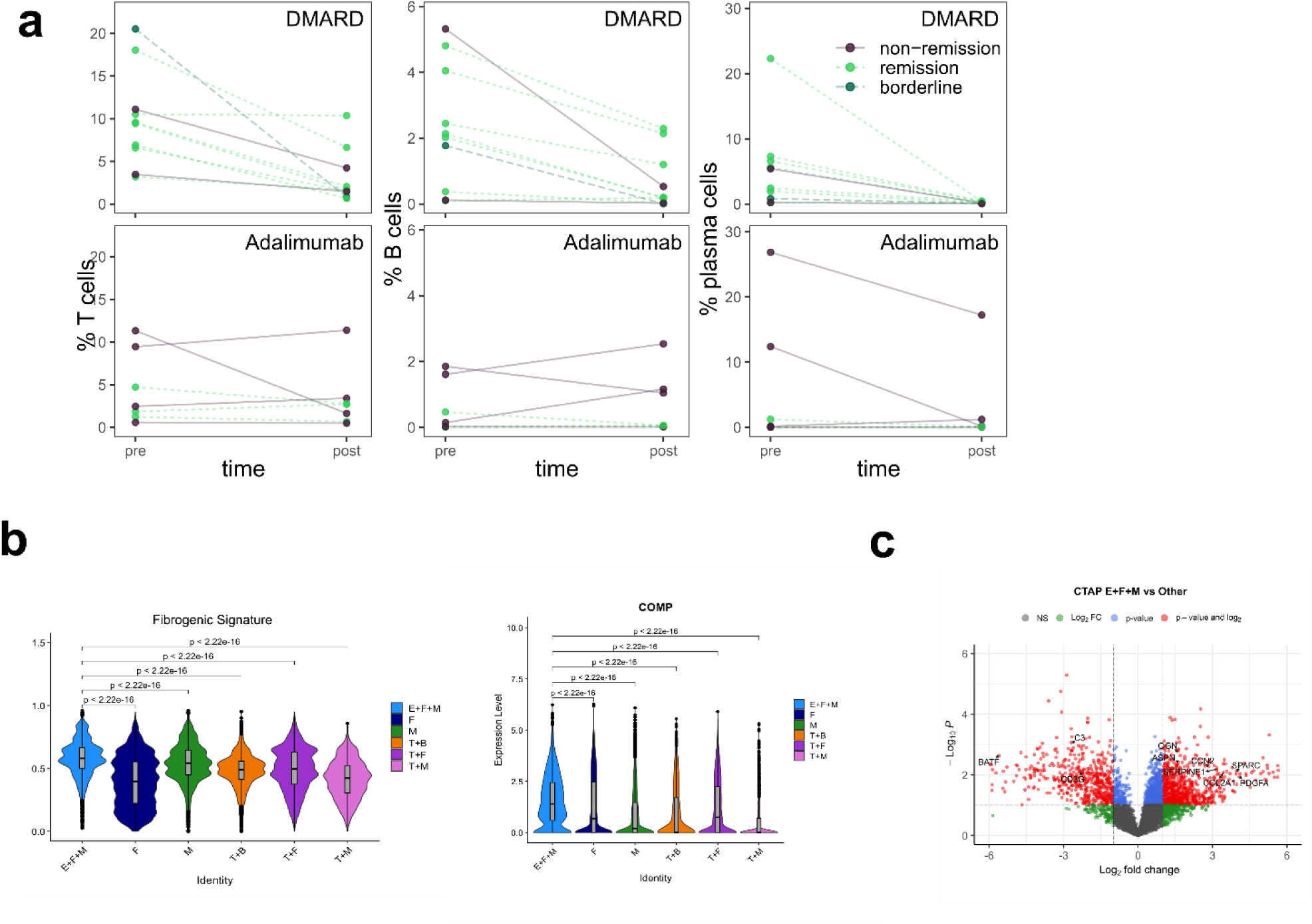
Pre- and post-treatment immune cell abundance and fibrogenic signature by CTAP. **a,** Quantification of the relative abundance of immune cell types pre- and post-treatment, separated by treatment type. **b,** Violin plot representing the distribution of fibrogenic gene signature score and *COMP* expression of synovial fibroblasts from each CTAP. Wilcoxon test was used for comparison between groups. **c,** Volcano plot of differentially abundant serum proteins in CTAP-EFM patients compared to patients assigned to other CTAPs with selected genes highlighted. P-values are unadjusted and were calculated using the limma package.

**Supplementary Table 1:** Clinical characteristics of patients enrolled in 396.10.

|  | **Total** | **remission** | **non-remission** | **borderline** |
| --- | --- | --- | --- | --- |
|  | (N=17) | (N=10) | (N=6) | (N=1) |
| **Age (y)** |  |  |  |  |
| Mean (SD) | 47.4 (17.4) | 50.3 (22.1) | 44.8 (5.19) | 33.0 (NA) |
| Median [Min, Max] | 45.0 [21.0, 75.0] | 56.5 [21.0, 75.0] | 44.5 [38.0, 54.0] | 33.0 [33.0, 33.0] |
| **Sex** |  |  |  |  |
| Female | 8 (47.1%) | 4 (40.0%) | 3 (50.0%) | 1 (100%) |
| Male | 9 (52.9%) | 6 (60.0%) | 3 (50.0%) | 0 (0%) |
| **DAS28-CRP (baseline)** |  |  |  |  |
| Mean (SD) | 4.90 (1.30) | 4.53 (1.34) | 5.28 (1.15) | 6.32 (NA) |
| Median [Min, Max] | 4.70 [2.50, 7.15] | 4.59 [2.50, 7.15] | 5.55 [3.77, 6.68] | 6.32 [6.32, 6.32] |
| **DAS28-ESR (baseline)** |  |  |  |  |
| Mean (SD) | 5.40 (1.54) | 4.89 (1.50) | 5.95 (1.39) | 7.31 (NA) |
| Median [Min, Max] | 5.21 [2.52, 8.06] | 5.08 [2.52, 7.79] | 6.12 [4.31, 8.06] | 7.31 [7.31, 7.31] |
| **Treatment** |  |  |  |  |
| DMARD | 10 (58.8%) | 7 (70.0%) | 2 (33.3%) | 1 (100%) |
| Adalimumab | 7 (41.2%) | 3 (30.0%) | 4 (66.7%) | 0 (0%) |

## Notes

### Competing Interest Statement

The authors have declared no competing interest.

